# Mifepristone decreases nicotine intake in dependent and non-dependent adult rats

**DOI:** 10.1101/2023.05.23.541914

**Authors:** Ranjithkumar Chellian, Azin Behnood-Rod, Adriaan W. Bruijnzeel

**Author notes:** Corresponding author: Adriaan Bruijnzeel, PhD University of Florida, Department of Psychiatry 1149 Newell Dr. Gainesville, Florida 32611.

## Abstract

Addiction to tobacco and nicotine products has adverse health effects and afflicts more than a billion people worldwide. Therefore, there is an urgent need for new treatments to reduce tobacco and nicotine use. Glucocorticoid receptor blockade shows promise as a novel treatment for drug abuse and stress-related disorders. The aim of these studies was to investigate if glucocorticoid receptor blockade with mifepristone diminishes the reinforcing properties of nicotine in rats with intermittent or daily long access to nicotine. The rats self-administered 0.06 mg/kg/inf of nicotine for 6 h per day, with either intermittent (3 days per week) or daily access (7 days per week) for 4 weeks before treatment with mifepristone. Daily nicotine self-administration models regular smoking, while intermittent nicotine self-administration models occasional smoking. To determine if the rats were dependent, they were treated with the nicotinic acetylcholine receptor antagonist mecamylamine, and somatic signs were recorded. The rats with intermittent access to nicotine had a higher level of nicotine intake per session than those with daily access, but only the rats with daily access to nicotine showed signs of dependence. Furthermore, mecamylamine increased nicotine intake during the first hour of access in rats with daily access but not in those with intermittent access. Mifepristone decreased total nicotine intake in rats with intermittent and daily access to nicotine. Moreover, mifepristone decreased the total distance traveled and rearing in the open field test and operant responding for food pellets. These findings indicate that mifepristone decreases the reinforcing effects of nicotine and food, but it might also be somewhat sedative.

## 1. Introduction

Tobacco addiction is a chronic disorder that is characterized by affective and somatic withdrawal signs, craving, and relapse after periods of abstinence [1]. Smoking increases the risk of more than a dozen types of cancer, cardiovascular disease, and Alzheimer’s disease [2–4]. Smoking is a major public health concern with about 1.3 billion smokers worldwide and 30 million in the United States. The great majority of smokers would like to quit, but only 3-5 percent are able to do so without medication [5]. The U.S. Food and Drug Administration has approved several smoking cessation treatments, but even when using these treatments, most people still relapse within a few months [6, 7]. Therefore, there remains an urgent need for new, efficacious, and safe smoking cessation treatments.

The glucocorticoid corticosterone orchestrates physiological and behavioral adaptations to stress. Glucocorticoids also have potent reinforcing properties. The synthetic cortisol derivative prednisone improves well-being and can induce euphoria in humans [8]. Moreover, rats prefer a corticosterone solution over tap water and self-administer glucocorticoids intravenously [9, 10]. Chronic corticosterone administration also facilitates responding in the intracranial self-stimulation procedure, which suggests that corticosterone potentiates brain reward function [11]. Moreover, glucocorticoids affect drug intake. Adrenalectomy decreases the acquisition of drug intake, reduces drug self-administration, and prevents drug sensitization [12, 13]. Additionally, adrenalectomy or corticosterone synthesis inhibition prevents stress-induced sensitization of the motor effects of psychostimulants [14, 15]. The glucocorticoid (Ki 0.09 nM) and progesterone (Ki 1 nM) receptor antagonist mifepristone (also called RU 486 or RU 38486) has been used to investigate the role of glucocorticoids in drug intake [16–20]. Treatment with mifepristone decreases the self-administration of cocaine and amphetamine in rats [21, 22]. The effects of mifepristone on the reinforcing effects of drugs in dependent animals have mainly been studied in the context of alcohol use. Mifepristone reduces alcohol intake in alcohol-dependent rats but does not affect alcohol intake in non-dependent animals with low levels of alcohol intake [20, 23]. Importantly, mifepristone decreases alcohol consumption and craving in people with alcohol use disorder [20]. Several studies have shown that selective glucocorticoid receptor antagonists have similar effects on alcohol intake as mifepristone, which suggests that the effects of mifepristone on drug intake are mediated via glucocorticoid, and not progesterone, receptor blockade [20, 24].

Clinical studies provide support for the role of stress and glucocorticoids in smoking [25, 26]. Stressors make it more difficult to resist smoking and lead to more intense smoking [25]. There is also evidence for the role of brain stress systems in the reinstatement of nicotine-seeking and nicotine withdrawal. Stressors induce the reinstatement of extinguished nicotine-seeking in rats [27]. Furthermore, cessation of nicotine administration leads to the activation of central stress systems, and this contributes to the anhedonia associated with nicotine withdrawal [28]. However, very little research has been done to investigate the role of stress systems in the self-administration of nicotine. A recent study showed that a pharmacological stressor increased the self-administration of nicotine in rats with a low level of nicotine intake but not in rats with a high level of nicotine intake [29]. Further research into the link between stress systems and nicotine self-administration could provide insight into the neurobiological mechanisms underlying nicotine dependence and contribute to the development of new smoking cessation treatments.

Studies with alcohol-dependent subjects suggest that glucocorticoid receptor blockade with mifepristone could be a novel approach for treating drug abuse [20]. However, it is currently unknown whether mifepristone affects smoking in humans or nicotine self-administration in rodents. To determine if mifepristone could help people quit smoking, we investigated the effect of mifepristone on nicotine self-administration in rats. The effects of mifepristone were investigated in rats with daily long access to nicotine and in rats with intermittent long access to nicotine. Daily nicotine self-administration models the smoking pattern in regular smokers, while intermittent nicotine self-administration models the sporadic smoking pattern in occasional smokers. Intermittent smokers are less dependent than regular smokers but still often have difficulty quitting smoking [30, 31]. Furthermore, we investigated the effects of mifepristone on operant responding for food and locomotor activity in the small open field test. Our findings indicate that mifepristone decreases the self-administration of nicotine in rats with both daily and intermittent long access to nicotine. This suggests that mifepristone might be an effective smoking cessation treatment for nicotine-dependent heavy smokers and for intermittent smokers who struggle to quit.

## 2. Materials and methods

### 2.1. Animals

Adult male Wistar rats (200 - 250 g, 8-9 weeks of age) were purchased from Charles River (Raleigh, NC). The rats were pair-housed in a climate-controlled vivarium on a reversed 12 h light-dark cycle (light off at 7 AM). The rats were gently handled for 2-3 min per day for several days before the start of the food training sessions. During the food training period, the rats were singly housed and remained so for the remainder of the study. Prior to the onset of the studies, food was available ad-lib in the home cage. During the food training and self-administration sessions, the rats were fed 90-95 percent of their ad-lib intake. Water was available ad-lib throughout the study. The experimental protocols were approved by the University of Florida Institutional Animal Care and Use Committee (IACUC). All experiments were performed in accordance with relevant guidelines and regulations of IACUC and in compliance with the ARRIVE guidelines 2.0 (Animal Research: Reporting of In Vivo Experiments).

### 2.2. Drugs

(-)-Nicotine hydrogen tartrate (Sigma-Aldrich, St. Louis, MO) was dissolved in sterile saline (0.9 % sodium chloride), and the pH was adjusted to 7.2 ± 0.2 using 1 M NaOH. The rats self-administered 0.03 or 0.06 mg/kg/inf of nicotine in a volume of 0.1 ml/inf. Mifepristone (Cayman chemical company, Ann Arbor, MI) was dissolved in 10 % dimethylformamide (Sigma-Aldrich, St. Louis, MO), 10 % kolliphor (Sigma-Aldrich, St. Louis, MO) and mixed in sterile saline. (-)-Nicotine hydrogen tartrate and mecamylamine hydrochloride (Sigma-Aldrich, St. Louis, MO) were dissolved in sterile saline. Mifepristone was administered intraperitoneally (IP), and mecamylamine and nicotine were administered subcutaneously (SC) in a volume of 1 ml/kg body weight. Nicotine doses are expressed as the base, and mecamylamine doses are expressed as salt.

### 2.3. Food training

A schematic overview of the experimental design is depicted in Figure 1. Rats were trained to press a lever for food pellets in operant chambers that were placed in sound- and light-attenuated cubicles (Med Associates, St. Albans, VT). Food training was conducted before the catheters were implanted. Responding on the active lever resulted in the delivery of a food pellet (45 mg, F0299, Bio-Serv, Frenchtown, NJ), and responding on the inactive lever was recorded but did not have scheduled consequences. Food delivery was paired with a cue light, which remained illuminated throughout the time-out (TO) period. The food training sessions were conducted for 10 days. Instrumental training started under an fixed-ratio 1, time-out 1s (FR1-TO1s) reinforcement schedule, and the rats remained on this schedule for 5 days (30 min sessions per day). After the fifth food training session, the rats were single-housed and remained so for the rest of the study. On day 6, the time-out period was increased to 10 s. The rats were allowed to respond for food pellets under the FR1-TO10s schedule (20 min sessions) for 5 days. Both levers were retracted during the 10 s time-out period. The rats were fed 90-95 percent of their baseline food intake in the home cage when the food training was ongoing.

**Figure 1.**
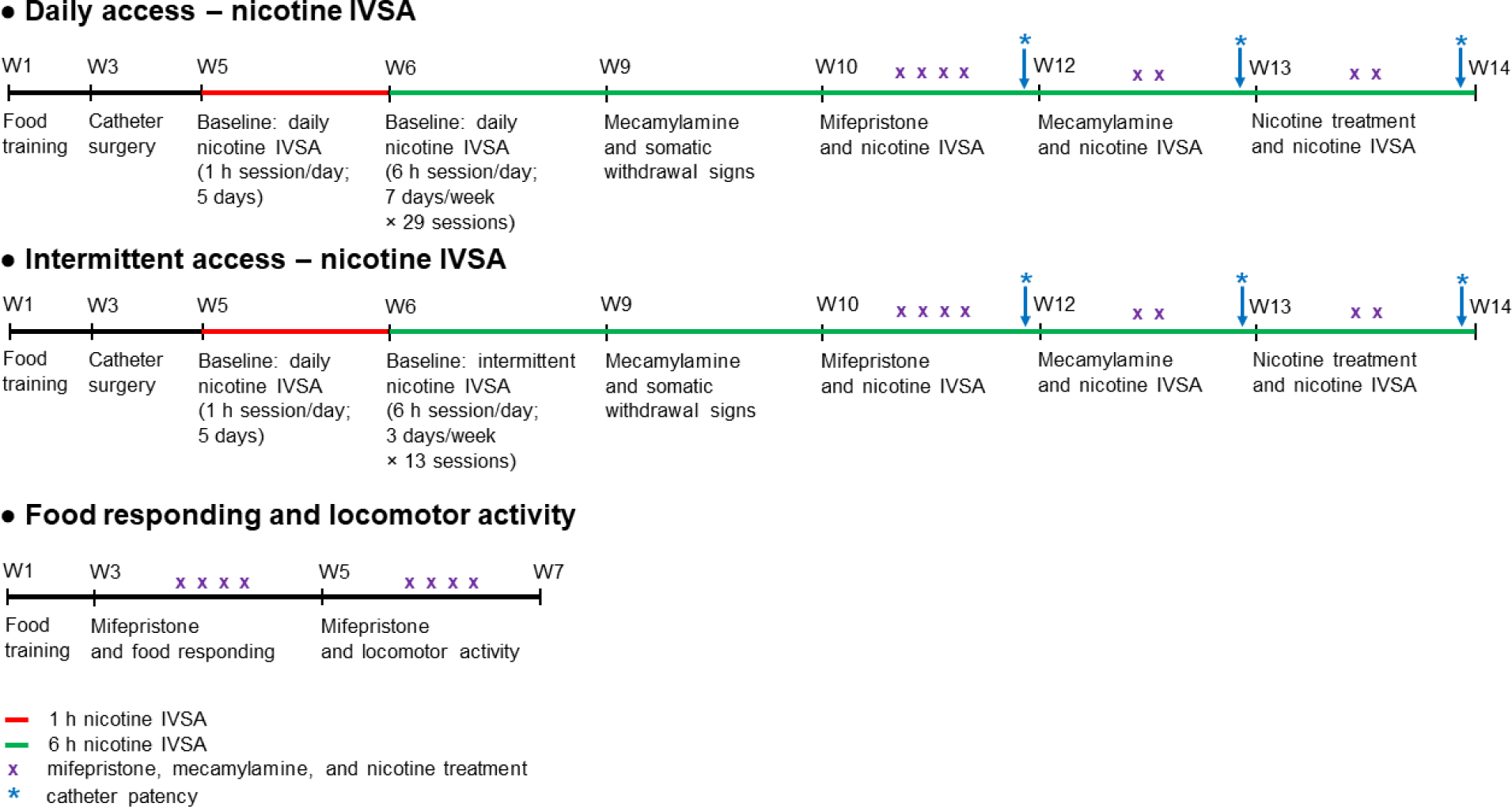
Schematic overview of the experiment. In experiment 1, rats were trained to respond for food pellets and were then prepared with IV catheters. The rats were then allowed to self-administer nicotine in five 1 h sessions and then either in 29 daily sessions or 13 intermittent sessions. After these sessions, the effects of mifepristone on nicotine self-administration were investigated. Subsequently, the effects of mecamylamine and nicotine pre-treatment on nicotine self-administration were investigated. Catheter patency was assessed following the completion of the mifepristone, mecamylamine, and nicotine treatments. In experiment 2, the rats were trained to respond for food pellets and then the effects of mifepristone on operant responding for food was investigated, and subsequently, the effects of mifepristone on locomotor activity, rearing, and stereotypies in the small open field were investigated. Abbreviations: W, weeks; IVSA, intravenous self-administration.

### 2.4. Intravenous catheter implantation

The catheters were implanted as described before [32–34]. The rats were anesthetized with an isoflurane-oxygen vapor mixture (1-3%) and prepared with a catheter in the right jugular vein. The catheters consisted of polyurethane tubing (length 10 cm, inner diameter 0.64 mm, outer diameter 1.0 mm, model 3Fr, Instech Laboratories, Plymouth Meeting, PA). The right jugular vein was isolated, and the catheter was inserted 2.9 cm. The tubing was then tunneled subcutaneously and connected to a vascular access button (Instech Laboratories, Plymouth Meeting, PA). The button was exteriorized through a 1-cm incision between the scapulae. During the 7-day recovery period, the rats received daily infusions of the antibiotic Gentamycin (4 mg/kg, IV, Sigma-Aldrich, St. Louis, MO). A sterile heparin solution (0.1 ml, 50 U/ml) was flushed through the catheter before and after administering the antibiotic or nicotine self-administration. After flushing the catheter or a nicotine self-administration session, 0.05 ml of a sterile heparin/glycerol lock solution (500 U/ml; Instech Laboratories, Plymouth Meeting, PA) was infused into the catheter. The animals received carprofen (5 mg/kg, SC; Patterson Veterinary, Loveland, CO) daily for 72 hours after the surgery. Two days before the start of the nicotine self-administration sessions, the rats were allowed to respond for food pellets under the FR1-TO10s schedule (one 20-min session).

### 2.5. Small open-field test

The small open field test was conducted as described before [35, 36]. The small open field test was conducted to assess locomotor activity, rearing, and stereotypies. These behaviors were measured using an automated animal activity cage system (VersaMax Animal Activity Monitoring System, AccuScan Instruments, Columbus, OH, USA). Horizontal beam breaks and total distance traveled reflect locomotor activity, and vertical beam breaks reflect rearing. The distance traveled is dependent on the path of the animal in the open field and better reflects locomotor activity than horizontal beam breaks. Repeated interruptions of the same beam are a measure of stereotypies (stereotypy count)[37]. The open field setup consists of four animal activity cages made of clear acrylic (40 cm × 40 cm × 30 cm; L x W x H), with 16 equally spaced (2.5 cm) infrared beams across the length and width of the cage. The beams were located 2 cm above the cage floor (horizontal activity beams). An additional set of 16 infrared beams were located 14 cm above the cage floor (vertical activity beams). All beams were connected to a VersaMax analyzer, which sent information to a computer that displayed beam data through Windows-based software (VersaDat software). The small open field test was conducted in a dark room, and the cages were cleaned with a Nolvasan solution (chlorhexidine diacetate) between animals. At the beginning of each test, the rat was placed in the center of the small open field, and activity was measured for 20 min.

### 2.6. Experimental design

#### 2.6.1. Experiment 1: effects of mifepristone, mecamylamine, and nicotine on operant responding for nicotine in rats with daily and intermittent access to nicotine

The rats were allowed to self-administer nicotine for five daily 1 h baseline sessions. During the first three sessions (days 1-3), the rats self-administered 0.03 mg/kg/inf of nicotine under an FR1-TO10s schedule. During the following two sessions (days 4 and 5), the rats self-administered 0.06 mg/kg/inf of nicotine under an FR1-TO60s schedule. Rats that self-administer 0.06 mg/kg/inf of nicotine have a higher level of nicotine intake and plasma nicotine levels and more somatic withdrawal signs compared to rats self-administering 0.03 mg/kg/inf [38–40]. High doses of nicotine can cause seizures in rodents [41–43]. To prevent seizures, the time-out period was increased from 10 to 60 s when the dose of nicotine was increased from 0.03 to 0.06 mg/kg/inf. Total nicotine intake over a 1 h nicotine self-administration period is not affected by the time-out period (10 – 60 s)[41]. During the first day that the rats received the 0.03 or 0.06 mg/kg/inf dose, nicotine intake was limited to prevent aversive effects (i.e., seizures and dysphoria). The maximum number of infusions was set to 20 on the first day that the rats received the 0.03 mg/kg/inf dose and to 10 on the first day that the rats received the 0.06 mg/kg/inf dose. After five days of nicotine self-administration, rats continued to self-administering nicotine under either a daily access (7 days/week) schedule for 29 sessions or an intermittent access (3 days/week; Monday, Wednesday, and Friday) schedule for 13 sessions. The rats with daily and intermittent access self-administered 0.06 mg/kg/inf nicotine under an FR1-TO60s schedule in 6 h sessions. Responding on the active lever resulted in the delivery of a nicotine infusion (0.1 ml infused over a 5.6-s period). The initiation of the delivery of an infusion was paired with a cue light, which remained illuminated throughout the time-out period. Responding on the inactive lever was recorded but did not have scheduled consequences. The active and inactive levers were retracted during the time-out period.

Mecamylamine-precipitated somatic withdrawal signs were recorded in rats with daily and intermittent access to nicotine. Somatic signs were observed in a transparent Plexiglas observation chamber (25 cm x 25 cm x 46 cm) with 1 cm of corncob bedding. The rats were habituated to the observation chambers for 5 min per day on 3 consecutive days. The daily access rats received saline injections immediately after the 22^nd^ nicotine self-administration session and received mecamylamine injections (2 mg/kg, SC) immediately after the 23^rd^ nicotine self-administration session. The rats with intermittent access received saline injections immediately after the 10^th^ nicotine self-administration session and received mecamylamine injections (2 mg/kg, SC) immediately after the 11 nicotine self-administration session. Ten minutes after the saline or mecamylamine injections, the rats were placed in the observation chamber, and somatic withdrawal signs were recorded for 10 min. The following somatic withdrawal signs were recorded: body shakes, head shakes, chews, teeth chattering, cheek tremors, gasps, writhes, ptosis, genital licks, foot licks, yawns, and escape attempts. Ptosis was counted once per minute if present continuously. Somatic signs were observed in a quiet, brightly lit room. The total number of somatic signs was the sum of the individual occurrences.

After 29 daily sessions or 13 intermittent sessions, the effects of mifepristone on nicotine self-administration (0.06 mg/kg/inf; FR1-TO60s schedule, 6 h) in rats with daily and intermittent access schedules were investigated (Expt. 1A, Fig. 1). Mifepristone (0, 10, 30, and 60 mg/kg, IP) was administered according to a Latin square design, 90 min before the nicotine self-administration sessions. There were at least 72 h between injections with mifepristone. The doses of mifepristone were based on previous studies with rats [20, 44]. After completing the mifepristone study, the rats continued to self-administer nicotine on the same daily and intermittent access schedule. Both groups of rats, had a 6 h self-administration session 72 h after the last mifepristone injection. In subsequent sessions, the effects of mecamylamine on nicotine self-administration (0.06 mg/kg/inf; FR1-TO60s schedule, 6 h) were investigated (Expt. 1B). Mecamylamine (0, 2 mg/kg, SC) was administered according to a Latin square design 10 min before the nicotine self-administration sessions. Seventy-two hours after the last mecamylamine treatment, the effects of nicotine treatment on nicotine self-administration (0.06 mg/kg/inf; FR1-TO60s schedule, 6 h) in rats with daily and intermittent access schedules were investigated (Expt. 1C). Nicotine (0, 0.4 mg/kg, SC) was administered according to a Latin square design 10 min before the nicotine self-administration sessions. It was also determined if mifepristone, mecamylamine, and nicotine affect operant responding for nicotine 24 and 48 h after treatment in the daily access rats. During the self-administration period, the rats received 90-95 percent of their normal ad-lib food intake in the home cage. The rats were fed (23 g) immediately after the operant sessions. A mild level of food restriction facilitates food training and nicotine self-administration in rats [45, 46]. During the 6 h nicotine self-administration sessions, the rats had access to water in the operant chambers. Catheter patency was assessed following the completion of the mifepristone, mecamylamine, and nicotine treatments (Fig. 1). The catheters were tested by infusing 0.2 ml of the ultra-short action barbiturate Brevital (1 % methohexital sodium). Rats with patent catheters displayed a sudden loss of muscle tone. If the rats did not respond to Brevital, their self-administration data were excluded from the analysis. Three rats from the daily access group did not respond to Brevital after completing the mecamylamine treatments. Therefore, the data from these rats were not included in the experiments that investigated the effects of mecamylamine and nicotine treatment on nicotine self-administration.

#### 2.6.2. Experiment 2: effects of mifepristone on operant responding for food and motor activity

After the food training sessions, the effects of mifepristone on operant responding for food pellets were investigated in daily 20-min sessions under an FR1-TO10s schedule (Expt. 2A). The rats were fed 90-95 percent of their baseline food intake in the home cage after operant responding for food. After the food study, the rats were fed ad-lib, and the effects of mifepristone on activity parameters in the small open field were investigated (Expt. 2B). The rats were habituated to the small open field on three consecutive days (20-min sessions), and then the effects of mifepristone on locomotor activity, rearing, and stereotypies in the small open field were investigated in 20-min sessions. In both experiments, mifepristone (0, 10, 30, and 60 mg/kg, IP) was administered according to a Latin square design 90 min before the food responding session or the small open field test. There were at least 72 h between injections with mifepristone. It was also determined if mifepristone affects operant responding for food and locomotor activity 24 and 48 h after treatment.

### 2.7. Statistics

Baseline nicotine self-administration data were analyzed using one-, two-, or three-way ANOVAs, with time (days), time point (hours), and session (first and last) as within-subjects factors and access schedule (daily and intermittent) and lever (active and inactive) as between-subjects factors (Table S1). Somatic withdrawal scores were analyzed with a two-way ANOVA, with mecamylamine treatment as a within-subjects factor and access schedule as a between-subjects factor. The effects of mifepristone, mecamylamine, and nicotine treatments on nicotine self-administration in rats with daily and intermittent access were analyzed using two- or three-way ANOVAs, with drug treatment and time point as within-subjects factors and access schedule as a between-subjects factor. The effects of mifepristone on operant responding on the active and inactive levers, food pellets received, and open-field behavior were analyzed with one-way ANOVAs, with drug treatment as a within-subjects factor. For all statistical analyses, significant effects in the ANOVA were followed by Bonferroni’s post hoc tests to determine which groups differed. P-values less than or equal to 0.05 were considered significant. Data were analyzed with SPSS Statistics version 29 and GraphPad Prism version 9.3.1. Figures and heatmaps were generated with GraphPad Prism version 9.3.1.

## 1.3. Results

### 3.1. Experiment 1: effects of mifepristone, mecamylamine, and nicotine on operant responding for nicotine in rats with daily and intermittent access to nicotine

#### 3.1.1. Baseline operant responding for nicotine

##### Nicotine 0.03 mg/kg/inf, three 1 h sessions

Nicotine intake (Fig. S1A) and responding on the active lever (Fig. S1B) decreased during the first three nicotine self-administration sessions (0.03 mg/kg/inf), and there was no difference in these parameters between the experimental groups (Nicotine intake: Time F2,48 =9.98, P < 0.001; Access F1,24=0.002, NS; Time x Access F2,48=1.252, NS; Active lever: Time F2,48 =10.071, P < 0.001; Access F1,24=0.002, NS; Time x Access F2,48=1.249, NS). Responding on the inactive lever (Fig. S1B) increased over time and was not affected by the experimental group (Inactive lever: Time F2,48 =8.199, P < 0.001; Access F1,24=1.217, NS; Time x Access F2,48=0.772, NS). **Nicotine 0.06 mg/kg/inf, two 1 h sessions:** The rats had access to 0.06 mg/kg/inf of nicotine (1 h/day) during the next two sessions. Nicotine intake, responding on the active lever, and responding on the inactive lever did not change over time and were not affected by the experimental group (Fig. S1C, Nicotine intake: Time F1,24 =1.782, NS; Access F1,24=0.17, NS; Time x Access F1,24=0.002, NS; Fig. S1D, Active lever: Time F1,24 =1.024, NS; Access F1,24=0.205, NS; Time x Access F1,24=0, NS; Fig. S1D, Inactive lever: Time F1,24 =0.716, NS; Access F1,24=0.281, NS; Time x Access F1,24=0.011, NS). **Nicotine 0.06 mg/kg/inf, twenty-nine 6 h sessions (daily, sessions 1-29):** The rats with daily long access self-administered nicotine for 29 session. During the 29 sessions (0.06 mg/kg/inf, 6 h/day), the daily long access rats pressed more on the active lever than on the inactive lever and lever pressing decreased over time (Fig. S2A, Active and inactive lever: Time F28,672 = 4.287, P < 0.001; Lever F1,24= 44.993, P < 0.001; Time x Lever F28,672 = 1.442, NS). During the 29 sessions, nicotine intake initially declined and reached a relatively stable level from day 7 onward (Fig. 2A, Time F28,336 = 5.066, P < 0.001). **Nicotine 0.06 mg/kg/inf, thirteen 6 h sessions (intermittent, sessions 1-13):** The rats with intermittent long access self-administered nicotine for 13 session. During the 13 sessions (0.06 mg/kg/inf, 6 h/day), the rats with intermittent long access responded more on the active lever than on the inactive lever and responding on the active lever was stable and responding on the inactive lever decreased (Fig. S2B, Active and inactive lever: Time F12,288 =1.156, NS; Lever F1,24=41.899, P < 0.001; Time x Lever F12,288 =2.547, P < 0.01). Nicotine intake did not change over time (Fig. 2B, Nicotine intake: Time F12,144 =1.717, NS).

**Figure 2.**
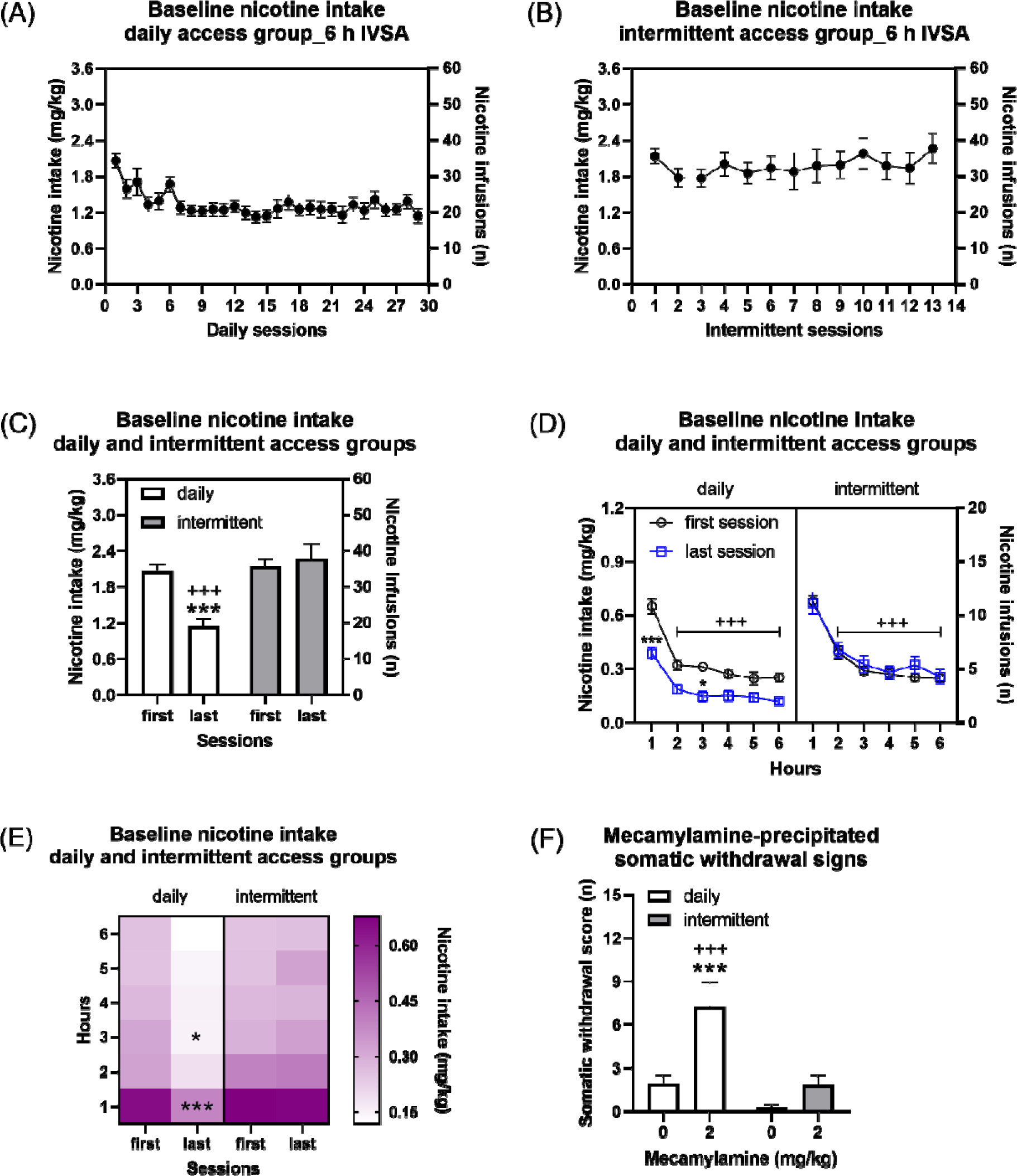
Baseline nicotine intake in rats with daily and intermittent access. The rats with daily access self-administered nicotine for 29 sessions and the rats with intermittent access self-administered nicotine for 13 sessions (A, daily; B, intermittent). All the sessions lasted 6 h and the rats self-administered 0.06 mg/kg/inf of nicotine. Nicotine intake was the same during the first and the last session in rats with intermittent access, but nicotine intake was lower during the last session in rats with daily access (C-E). Rats with daily, but not intermittent access to nicotine displayed somatic withdrawal signs (F). Asterisks indicate lower nicotine intake during the last than the first session in rats with daily access (C-E), and more somatic signs after treatment with mecamylamine than vehicle in rats with daily access to nicotine (F). Plus signs indicate lower nicotine intake compared to rats with intermittent access during the last session (C), lower nicotine intake compared to the first hour of nicotine intake during the first and last session in rats with the same access schedule (D), and more somatic signs compared to rats with intermittent access treated with mecamylamine (F). * P<0.05; ***, +++ P<0.001. Daily n=13, Intermittent, n=13. Data are expressed as means ± SEM.

##### Nicotine 0.06 mg/kg/inf, thirteen 6 h sessions, daily compared to intermittent

An additional analysis was conducted to compare operant responding for nicotine between rats with daily and intermittent access during the first thirteen sessions (Fig. S2C and S2D). The rats with intermittent access to nicotine had a higher level of nicotine intake and responded more on the active lever than rats with daily access (Fig. S2C, Nicotine intake: Access F1,24=6.683, P < 0.05; Fig. S2D, Active lever: Access F1,24=7.307, P < 0.05). Furthermore, nicotine intake and responding on the active lever were stable in the rats with intermittent access but decreased in then stabilized in rats with daily access (Fig. S2C, Nicotine intake: Time F12,288=3.652, P<0.001; Time x Access F12,288=5.018, P < 0.01; Fig. S2D, Active lever: Time F12,288=3.362, P<0.001; Time x Access F12,288=4.983, P <0.001). The post hoc showed that after 7 sessions, nicotine intake was lower in the rats with daily access than in rats with intermittent access. Responding on the inactive lever decreased over time and was not affected by the access schedule (Fig. S2D, Inactive lever: Access F1,24=0.01, NS; Time F12,288=2.491, P<0.01; Time x Access F12,288=0.819, NS).

##### Nicotine 0.06 mg/kg/inf, 6 h sessions, daily compared to intermittent (first and last session)

The rats with intermittent access to nicotine had a higher level of nicotine intake and responded more on the active lever than those with daily access (Fig. 2C and Fig. S2E, Nicotine intake: Access F1,24=10.283, P < 0.01; Active lever: Access F1,24=10.083, P < 0.01). The posthoc tests showed that nicotine intake and active lever responses were higher in rats with intermittent access than in rats with daily access during the last session (Fig. 2C and Fig. S2E). Nicotine intake and responding on the active lever decreased in the rats with daily access and did not change in the rats with intermittent access (Fig. 2C, Nicotine intake: Session F1,24 =8.934, P < 0.01; Session x Access F1,24 =15.698, P < 0.001; Fig. S2E, Active lever: Session F1,24 =10.08, P < 0.01; Session x Access F1,24 =15.365, P < 0.001). The posthoc analyses showed that nicotine intake and active lever responses were lower during the last session than during the first session in the rats with daily access to nicotine (Fig. 2C and Fig. S2E). There was no difference in responding on the inactive lever between the rats with daily and intermittent access (Fig. S2F, Inactive lever: Access F1,24=0.135, NS; Time F1,24 =3.038, NS; Time x Access F1,24 =0.05, NS).

##### Nicotine 0.06 mg/kg/inf, 6 h session, daily compared to intermittent (time course analyses of nicotine intake during first and last session)

An analysis was done to compare the time course (hourly intake and lever pressing over the 6 h period) of nicotine intake during the first and the last self-administration session in the rats with daily and intermittent access (Fig. 2D). The rats with intermittent access had a similar level of nicotine intake during the first and last session, but nicotine intake was lower during the last session than during the first session in the rats with daily access (Session F1,24= 8.934, P<0.01; Access F1,24= 10.283, P<0.01; Session x Access F1,24= 15.698, P<0.001). The posthoc test showed that nicotine intake was lower during the last session than during the first session in the rats with daily access at the 1 h and 3 h time points (Fig. 2D and 2E). Over the course of the 6 h self-administration period, nicotine intake decreased, and this decline was smallest during the last session in rats with daily access (Time points F5,120= 188.925, P<0.001; Time points x Access F5,120= 3.618, P<0.01; Time points x Session F5,120= 3.053, P<0.05; Time points x Access x Session F5,120= 1.129, NS).

#### 3.1.2. Somatic withdrawal signs (daily and intermittent groups)

Treatment with mecamylamine increased somatic withdrawal signs and the rats with daily access had more somatic withdrawal signs than the rats with intermittent access (Fig. 2F, Treatment F1,24 =22.993, P < 0.001; Access F1,24=9.042, P < 0.01; Treatment x Access F1,24=6.97, P < 0.05). The post hoc analysis revealed that rats with daily access to nicotine and treated with mecamylamine had significantly more somatic signs than those treated with saline. Moreover, mecamylamine induced more somatic withdrawal signs in rats with daily access to nicotine than in those with intermittent access to nicotine (Fig. 2F).

#### 3.1.3. Experiment 1A: effects of mifepristone on operant responding for nicotine

##### 90-min after mifepristone treatment (daily and intermittent groups)

Mifepristone induced a decrease in nicotine intake and responding on the active lever in rats with daily and intermittent access to nicotine (Fig. 3A, Table S2, Nicotine intake: Treatment F3,72 =27.398, P < 0.001; Treatment x Access F3,72 =5.704, P < 0.01; Fig. 3B, Active lever: Treatment F3,72 =27.737, P < 0.001; Treatment x Access F3,72 =5.624, P < 0.01). The posthoc test showed that 30 and 60 mg/kg of mifepristone decreased nicotine intake and responding on the active lever in daily access rats and 60 mg/kg of mifepristone decreased nicotine intake and responding on the active lever in intermittent access rats (Fig. 3A and 3B). The rats with intermittent access had a higher level of nicotine intake and responded more on the active lever than the rats with daily access (Fig 3A, Nicotine intake: Access F1,24=13.225, P < 0.01; Fig 3B, Active lever: Access F1,24=12.628, P < 0.01). There was no difference in responding on the inactive lever between the rats with daily and intermittent access and mifepristone did not affect responding on the inactive lever (Fig. 3C, Inactive lever: Treatment F3,72 =1.943, NS; Access F1,24=1.281, NS; Treatment x Access F3,72 =1.242, NS).

**Figure 3.**
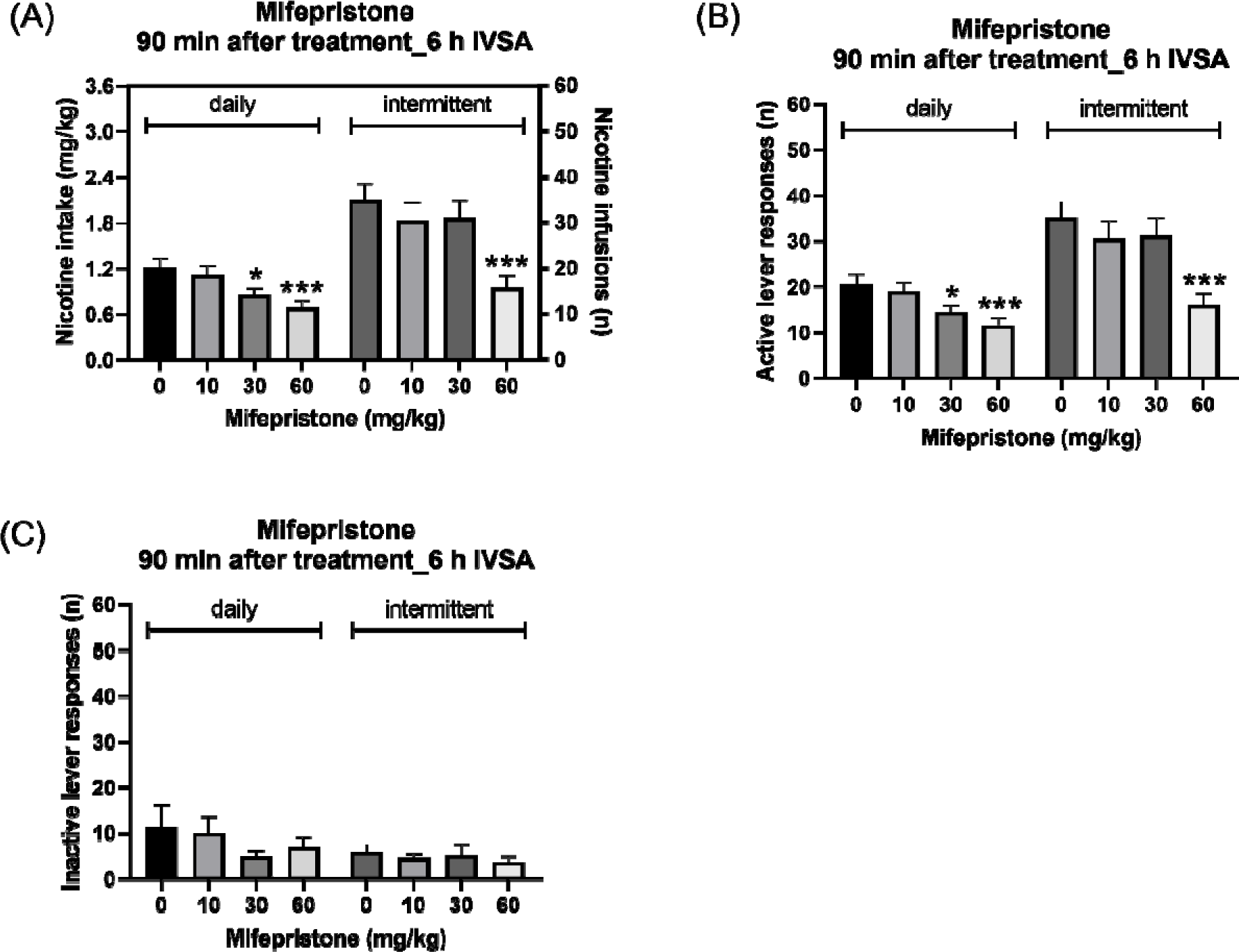
Mifepristone decreases nicotine intake. The effects of mifepristone on nicotine intake (A), active lever presses (B), and inactive lever presses (C) were investigated. Mifepristone decreased nicotine intake, responding on the active lever, but not responding on the inactive lever. Asterisks indicate lower nicotine intake and fewer lever presses compared to rats on the same access schedule and treated with vehicle. * P<0.05, *** P<0.001. Daily n=13, Intermittent, n=13. Data are expressed as means ± SEM.

##### Time course analysis, 90-min after mifepristone treatment (daily and intermittent groups)

A time course analysis was conducted to determine the duration of the effects of mifepristone in the rats with daily and intermittent access. The rats with intermittent access to nicotine had a higher level of nicotine intake during the 6 h self-administration session (Fig. 4A, 4B, and 4C, Access F1,24 =13.225, P < 0.01; Time points F5,120 =64.334, P < 0.001; Time points x Access F5,120 =8.121, P < 0.001). Mifepristone decreased nicotine intake in the rats with daily and intermittent access, with the greatest effect observed at the onset of the 6 h self-administration period (Fig. 4A, 4B, and 4C, Nicotine intake: Treatment F3,72 =27.398, P < 0.001; Time points x Treatment F15,360 =10.516, P < 0.001). Moreover, mifepristone induced a greater decrease in nicotine intake in rats with intermittent access to nicotine than in rats with daily access to nicotine (Fig. 4A, and 4B, Treatment x Access F3,72 =5.704, P < 0.01; Time points x Treatment x Access F15,360 =2.643, P < 0.001). The posthoc analysis showed that in rats with daily access to nicotine, 30 and 60 mg/kg of mifepristone decreased nicotine intake for 1 h (Fig. 4A, and 4C). Furthermore, the posthoc findings showed that 30 mg/kg of mifepristone decreased nicotine intake for 1 h, while a higher does, 60 mg/kg, decreased nicotine intake for 2 h in rats with intermittent access to nicotine (Fig. 4B and 4C).

**Figure 4.**
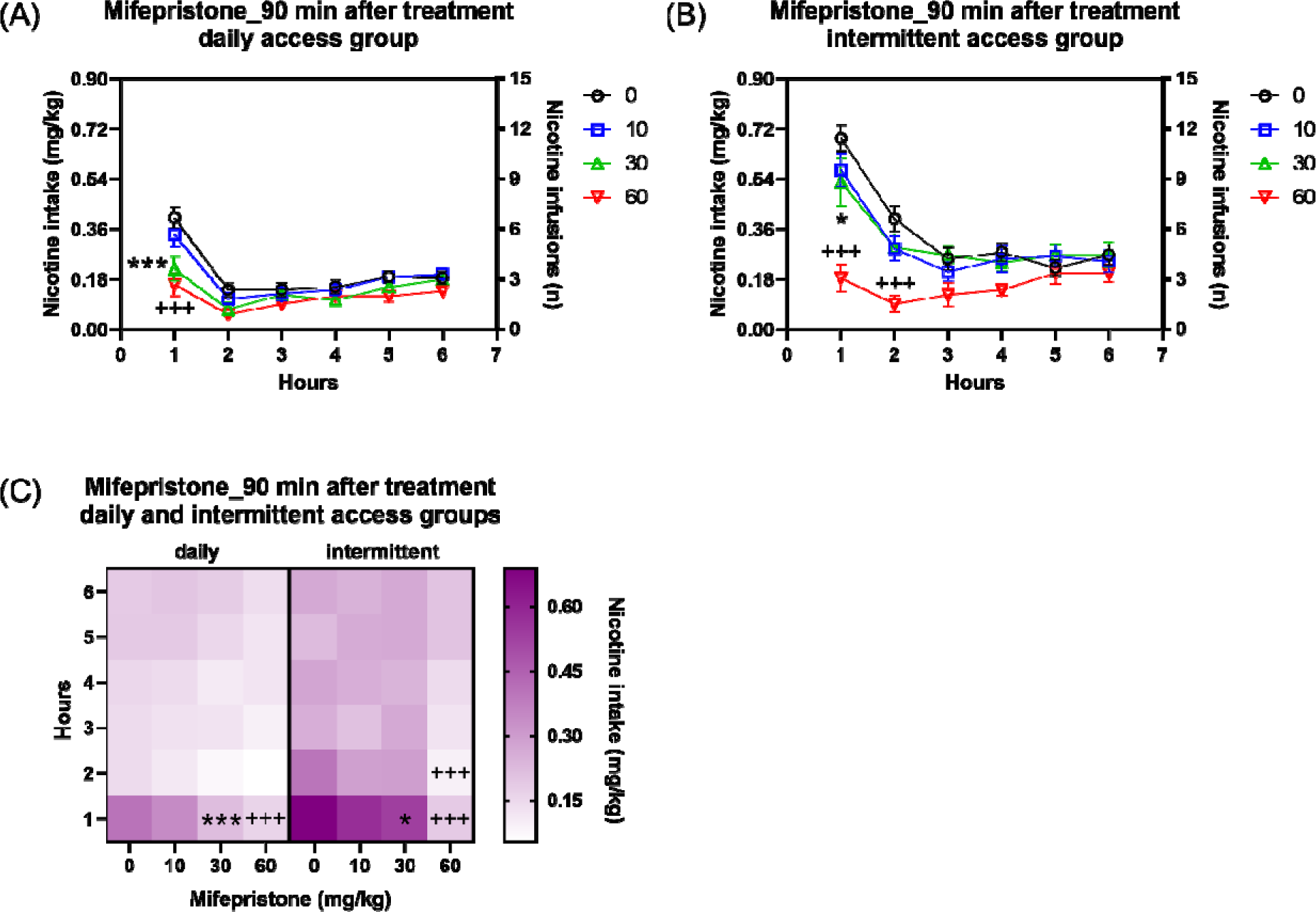
Time course effects of mifepristone on nicotine intake. Mifepristone decreased nicotine intake at the onset of the nicotine self-administration period in rats with daily (A) and intermittent (B) access. The heatmap shows the time course effects of mifepristone on nicotine intake in rats with daily and intermittent access (C). Asterisks indicate lower nicotine intake in rats treated with 30 mg/kg of mifepristone than in vehicle-treated rats with the same access schedule and at the same time point. Plus sign indicate lower nicotine intake in rats treated with 60 mg/kg of mifepristone than in vehicle-treated rats with the same access schedule and at the same time point. * P<0.05; ***, +++ P<0.001. Daily n=13, Intermittent, n=13. Data are expressed as means ± SEM.

##### 24 and 48 h after mifepristone treatment (daily group)

Mifepristone did not affect nicotine intake and pressing on the active lever or inactive lever 24 h and 48 h later (24 h time point; Fig. S3A, Nicotine intake: Treatment F3,36 =0.82, NS; Fig. S3B, Active lever: Treatment F3,36 =0.725, NS; Fig. S3C, Inactive lever: Treatment F3,36 =2.552, NS; 48 h time point; Fig. S3E, Nicotine intake: Treatment F3,36 =0.978, NS; Fig. S3F, Active lever: Treatment F3,36 =1.54, NS; Fig. S3G, Inactive lever: Treatment F3,36 =1.416, NS).

##### Time course analysis, 24 and 48 h after mifepristone treatment (daily group)

Mifepristone did not affect the time course of nicotine intake 24 h (Fig. S3D, Nicotine intake: Treatment F3,36 =0.82, NS; Time point F5,60 =63.045, P < 0.001; Treatment x Time point F15,180 =0.54, NS) or 48 h later (Fig. S3H, Nicotine intake: Treatment F3,36 =0.978, NS; Time point F5,60 =93.473, P < 0.001; Treatment x Time point F15,180 =1.024, NS).

#### 3.1.4. Experiment 1B: effects of mecamylamine on operant responding for nicotine

##### 10-min after mecamylamine treatment (daily and intermittent groups)

Treatment with mecamylamine did not affect nicotine intake or responding on the active lever over the 6 h self-administration period in rats with daily or intermittent access to nicotine (Fig. 5A, Nicotine intake: Treatment F1,21 =0.224, NS; Fig. S4A, Active lever: Treatment F1,21 =0.194, NS). The rats with intermittent access to nicotine self-administered more nicotine and had more active lever presses than rats with daily access (Fig. 5A, Nicotine intake: Treatment x Access F1,21 =3.024, NS; Access F1,21=9.202, P < 0.01; Fig. S4A, Active lever: Treatment x Access F1,21 =2.376, NS; Access F1,21=8.943, P < 0.01). Mecamylamine increased inactive lever presses in rats with daily and intermittent access, and there was no difference in inactive lever responses between rats with daily and intermittent access (Fig. S4B, Inactive lever: Treatment F1,21 =8.75, P < 0.01; Treatment x Access F1,21 =0.336, NS; Access F1,21=0.12, NS). The post hoc tests revealed that mecamylamine increased responding on the inactive lever in rats with intermittent access to nicotine (Fig. S4B).

**Figure 5.**
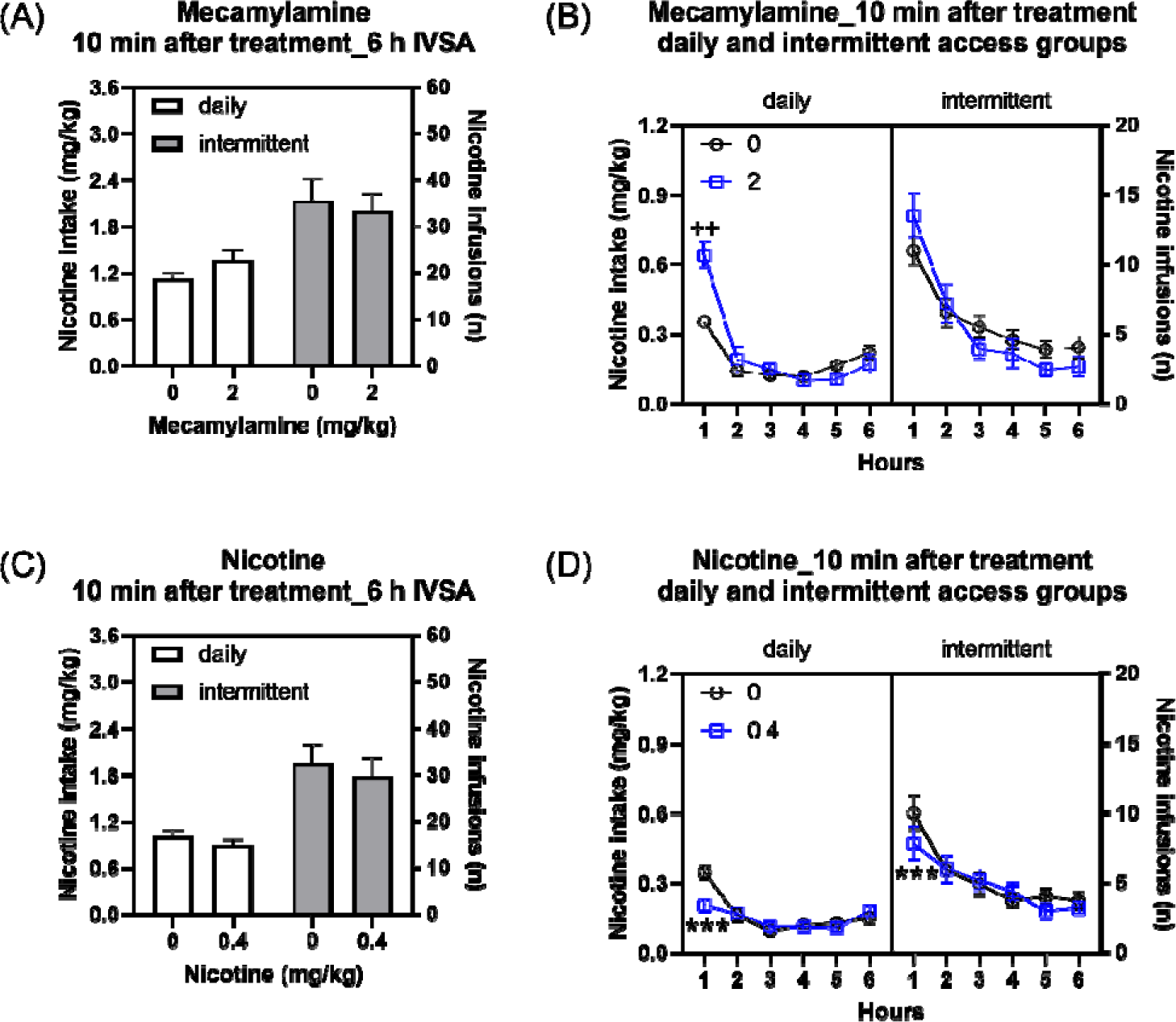
Effects of mecamylamine and nicotine pre-treatment on nicotine intake. The effects of mecamylamine (A, total intake; B, time-course) and nicotine (C, total intake; D, time-course) on nicotine intake was investigated. Mecamylamine did not affect total nicotine intake over the 6 h nicotine self-administration period in rats with daily and intermittent access to nicotine (A). Mecamylamine increased nicotine intake during the first hour of the self-administration session in the rats with daily access but not with intermittent access (B). Pre-treatment with nicotine decreased nicotine intake during the 6 h nicotine self-administration period in rats with daily and intermittent access to nicotine (C). Pre-treatment with nicotine also induced a decrease in nicotine intake during the first hour of the self-administration session in the rats with daily and intermittent access (D). Plus signs indicate higher nicotine intake in rats treated with 2 mg/kg of mecamylamine than in vehicle-treated rats with the same access schedule and at the same time point. Asterisks indicate lower nicotine intake in rats treated with 0.4 mg/kg of nicotine than in vehicle-treated rats with the same access schedule and at the same time point. ++ P<0.01; *** P<0.001. Daily n=10, Intermittent, n=13. Data are expressed as means ± SEM.

##### Time course analysis, 10-min after mecamylamine treatment (daily and intermittent groups)

The time course analysis showed that mecamylamine increased nicotine intake at the onset of the 6 h nicotine self-administration period only in rats with daily access to nicotine (Fig. 5B, Nicotine intake: Treatment F1,21 =0.224, NS; Treatment x Access F1,21 =3.024, NS; Treatment x Time point F5,105 =7.97, P < 0.001; Treatment x Time point x Access F5,105 =0.454, NS). The posthoc test showed that in rats with daily access, mecamylamine increased nicotine intake during the first hour of access, but not at later time points (Fig. 5B). The rats with intermittent access to nicotine had a higher level of nicotine intake than the rats with daily access during the 6 h self-administration session (Fig. 5B, Access F1,21 =9.202, P < 0.01; Time point F5,105 =75.473, P < 0.001; Time point x Access F5,105 =5.9, P < 0.001).

##### 24 and 48 h after mecamylamine treatment (daily group)

Mecamylamine did not affect nicotine intake, or active and inactive lever responses, 24 h and 48 h after treatment (see supplemental file for results; Fig. S5A-F and S6A-B).

#### 3.1.5. Experiment 1C: effects of nicotine pre-treatment on operant responding for nicotine

##### 10-min after nicotine treatment (daily and intermittent groups)

Treatment with nicotine decreased nicotine intake and responding on the active lever to a similar degree in the rats with daily and intermittent access to nicotine (Fig. 5C, Nicotine intake: Treatment F1,21 =7.303, P < 0.05; Treatment x Access F1,21 =0.223, NS; Access F1,21=10.486, P < 0.01; Fig. S4C, Active lever: Treatment F1,21 =5.114, P < 0.05; Treatment x Access F1,21 =0.001, NS; Access F1,21=11.311, P < 0.01). The post hoc test did not reveal any significant effects. Nicotine treatment did not affect responding on the inactive lever and there was no difference in inactive lever responses between the rats with daily and intermittent access (Fig. S4D, Inactive lever: Treatment F1,21 =0.081, NS; Treatment x Access F1,21 =0.013, NS; Access F1,21=0.124, NS).

##### Time course analysis, 10-min after nicotine treatment (daily and intermittent groups)

The time course analysis showed that pre-treatment with nicotine affected nicotine intake in rats with daily and intermittent access during the 6 h self-administration period (Fig. 5D, Nicotine intake: Treatment F1,21 =7.303, P < 0.05; Treatment x Access F1,21 =0.223, NS; Treatment x Time point F5,105 =10.129, P < 0.001; Treatment x Time point x Access F5,105 =0.973, NS). The posthoc analyses showed that nicotine pre-treatment decreased nicotine intake in rats with daily and intermittent access to nicotine during the first hour of the 6 h nicotine self-administration period (Fig. 5D). The rats with intermittent access to nicotine had a higher level of nicotine intake than the rats with daily access during the 6 h self-administration session (Fig. 5D, Access F1,21 =10.486, P < 0.01; Time point F5,105 =33.329, P < 0.001; Time point x Access F5,105 =6.402, P < 0.001).

##### 24 and 48 h after nicotine treatment (daily group)

Nicotine pre-treatment did not affect nicotine intake, or active and inactive lever responses, 24 h and 48 h after treatment (see supplemental file for results; Fig. S5G-L and S6C-D).

### 3.2. Experiment 2: effects of mifepristone on operant responding for food and motor activity

#### 3.2.1. Experiment 2A: effects of mifepristone on operant responding for food

##### 90-min after mifepristone treatment

Mifepristone decreased food intake and responding on both the active and inactive lever (Fig. 6A; Food intake: Treatment F3,27 =6.794, P < 0.001; Fig. 6B, Active lever: Treatment F3,27 =6.971, P < 0.001; Fig. 6C, Inactive lever: Treatment F3,27 =3.115, P < 0.05). The posthoc tests showed that 30 and 60 mg/kg of mifepristone decreased food intake and responding on the active lever and 30 mg/kg of mifepristone decreased responding on the inactive lever.

**Figure 6.**
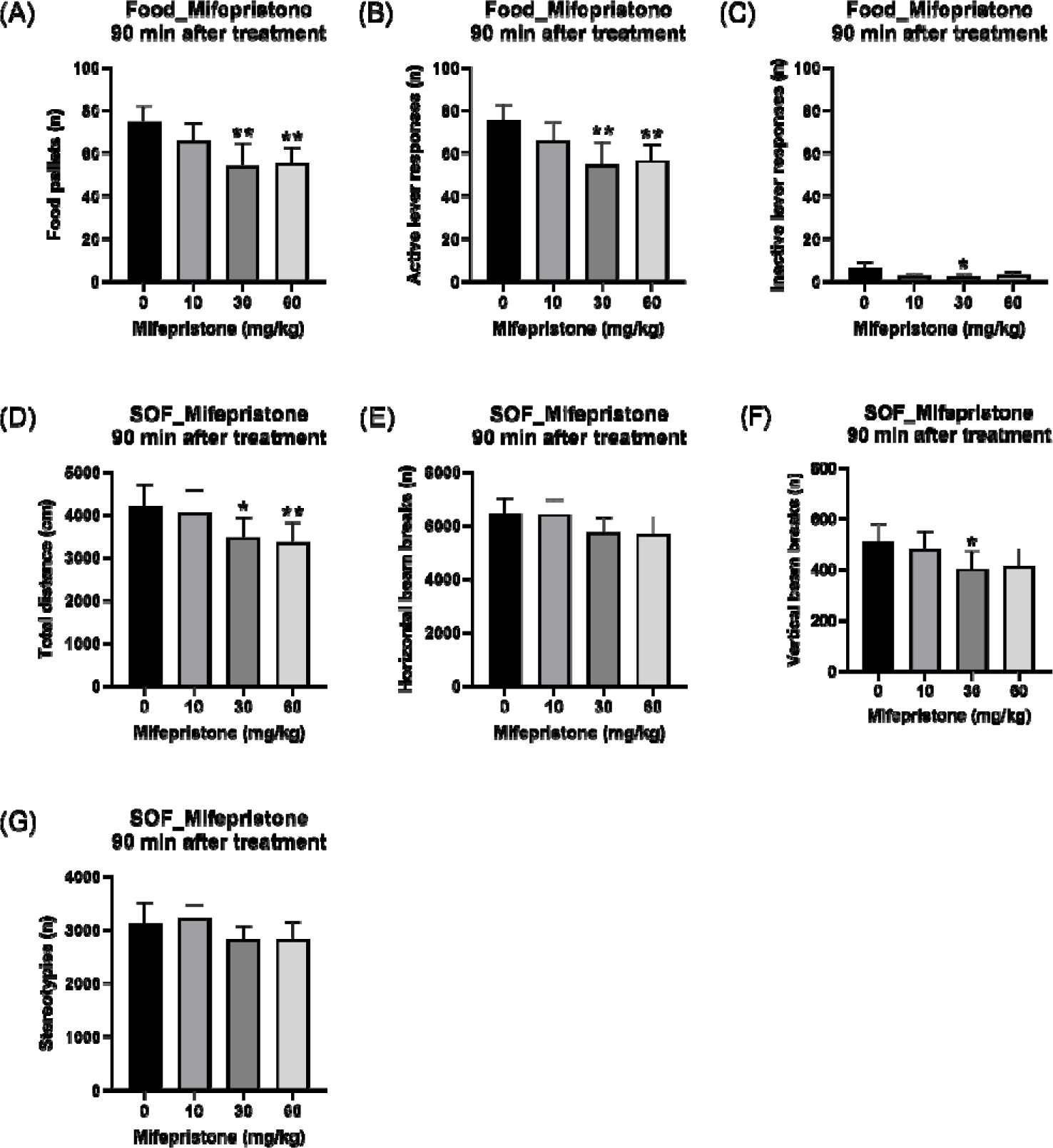
Mifepristone decreases operant responding for food, total distance traveled, and rearing. The effects of mifepristone on food pellets received (A), active lever presses (B), and inactive lever presses (C) was investigated. Furthermore, the effects of mifepristone on the total distance traveled (D), horizontal beam breaks (E), vertical beam breaks (F), and stereotypies (G) was investigated. Mifepristone decreased food intake, responding on the active lever, and responding on the inactive lever (A-C). Mifepristone also decreased the total distance traveled (D) and vertical beam breaks (F) and did not affect horizontal beam breaks (C) and stereotypies (G). N=10. Asterisks indicate significantly different compared to rats treated with vehicle. * P<0.05; ** P<0.01. Data are expressed as means ± SEM.

##### 24 and 48 h after mifepristone treatment

Mifepristone did not affect food intake, or active and inactive lever responses, 24 h and 48 h after treatment (see supplemental file for results; Fig. S7A-F).

#### 3.2.2. Experiment 2B: effects of mifepristone on locomotor activity

##### 90-min after mifepristone treatment

Treatment with mifepristone decreased the total distance traveled and vertical beam breaks (Fig. 6D, Total distance: Treatment F3,27 =5.704, P < 0.01; Fig. 6F, Vertical beam breaks: Treatment F3,27 =3.032, P < 0.05). The posthoc tests showed that 30 and 60 mg/kg of mifepristone decreased the total distance traveled and 30 mg/kg of mifepristone decreased vertical beam breaks (Fig. 6D and 6F). Mifepristone did not affect horizontal beam breaks and stereotypies (Fig. 6E, Horizontal beam breaks: Treatment F3,27 =2.861, P = 0.055; Fig. 6G, Stereotypies: Treatment F3,27 =1.904, NS).

##### 24 and 48 h after mifepristone treatment

Mifepristone did not affect total distance, horizontal and vertical beam breaks, and stereotypies 24 h and 48 h after treatment (see supplemental file for results; Fig. S8A-H).

## 1.4. Discussion

The current study explored the effects of the glucocorticoid and progesterone receptor antagonist mifepristone on nicotine intake in rats with daily and intermittent long access to nicotine. Rats with daily access to nicotine developed nicotine dependence, as indicated by an increase in somatic withdrawal signs after treatment with mecamylamine. In contrast, rats with intermittent access did not become dependent despite having a higher level of nicotine intake per session than the rats with daily access. Mifepristone induced a greater decrease in nicotine intake in rats with intermittent access than in rats with daily access. Moreover, mifepristone decreased the total distance traveled and rearing in the small open field test and operant responding for food. These findings demonstrate that the blockade of glucocorticoid receptors decreases operant responding for nicotine and food and may also have some sedative effects.

In this study, the rats were given access to a 0.03 mg/kg/inf dose of nicotine for three days and a 0.06 mg/kg/inf dose for two days (1 h sessions) prior to the initiation of the long access sessions. There was no difference in nicotine intake between the two groups during these initial five days. However, during the long access sessions, the rats with intermittent access to nicotine had a higher level of nicotine intake per session than rats with daily access. This outcome is in line with prior research that showed that rats with intermittent access have higher nicotine intake per session than those with daily access [47]. When the rats in the present study were given long access to nicotine, nicotine intake in the rats with daily access decreased over time, whereas nicotine intake in the rats with intermittent access remained unchanged. Cohen et al. found that male rats increased their nicotine intake when switched from daily long access to intermittent long access, but there was no further increase after they had been switched to intermittent access [47]. In a prior study by our group, we found a small increase in nicotine intake over time in male rats with intermittent long access to nicotine [48]. Overall, these findings indicate that rats with intermittent access have higher nicotine intake per session than those with daily access. However, neither intermittent nor daily access leads to a robust escalation of drug intake, as observed with cocaine [49].

In the current study, mecamylamine increased somatic withdrawal signs in rats with daily access to nicotine but not in those with intermittent access. This suggests that daily access to nicotine, but not intermittent access, leads to the development of dependence. This finding is in line with a previous study, which demonstrated that animals that self-administered nicotine for 7 days a week, but not 5 days a week, displayed withdrawal signs after mecamylamine treatment [40, 50]. This pattern of results demonstrates that animals with intermittent access to nicotine have higher nicotine intake per session compared to those with daily access. However, the rats with intermittent access do not become dependent as indicated by the absence of somatic withdrawal signs.

In this study, we investigated the effects of mifepristone on nicotine self-administration in rats with daily access (dependent) and intermittent access (non-dependent) to nicotine. Mifepristone decreased nicotine intake in both groups of rats. Mifepristone decreased nicotine intake during the first hour of the self-administration session in rats with daily access, while it reduced nicotine intake during the first two hours in rats with intermittent access. This suggests that mifepristone has a greater effect on nicotine intake in non-dependent rats than in nicotine-dependent rats. Mifepristone did not affect nicotine intake at the 24-h and 48-h time points in both nicotine-dependent and non-dependent rats, indicating that it does not have a long-term effect on nicotine intake. A previous study showed that mifepristone prevented the escalation of alcohol intake and the motivation for alcohol intake in alcohol-dependent rats. However, mifepristone did not affect alcohol intake in rats that were not alcohol-dependent [23]. Other studies also reported that acute treatment with mifepristone did not affect alcohol intake in non-dependent animals [51–54]. Therefore, these findings suggest that mifepristone differently affects alcohol and nicotine intake. Mifepristone decreased nicotine intake in dependent and non-dependent animals, while its effects on alcohol intake were only observed in dependent animals. In the present study, we investigated the effects of mifepristone on nicotine intake only in male Wistar rats. Recent studies suggest that there are no sex differences in the effects of mifepristone (≥ 30 mg/kg) on alcohol and saccharin self-administration, as well as anxiety-like behavior in Wistar rats [44, 55]. Thus, these findings indicate that there are no sex differences in the effects of mifepristone, and it is unlikely that mifepristone would have affected nicotine intake differently between male and female rats.

We also investigated the effects of mifepristone on operant responding for food pellets. Mifepristone reduced operant responding for food pellets, suggesting that mifepristone affects the reinforcing properties of non-drug rewards. The effects of mifepristone were observed 90 min after treatment but not at later time points, indicating that mifepristone does not have long-term effects on food intake. Consistent with our findings, a recent study showed that 60 mg/kg of mifepristone decreased saccharin self-administration in male and female Wistar rats [44]. In contrast, low doses (≤ 30 mg/kg) of mifepristone did not affect the self-administration of saccharin or sucrose solutions in male Wistar and Long-Evans rats [20, 56]. With regard to water self-administration, 30 mg/kg of mifepristone decreased water intake, while 60 mg/kg of mifepristone did not affect water intake in male Wistar rats [20]. Additionally, mifepristone doses that decreased alcohol intake in male and female HDID-1 mice (50 and 100 mg/kg) and rhesus monkeys (30 and 56 mg/kg) did not affect water intake [57]. Mifepristone also decreases food intake in people treated with the antipsychotic risperidone and olanzapine [58, 59]. These findings suggest that mifepristone decreases the reinforcing properties of food in rodents and humans.

The current study also examined whether mifepristone affects behavior in the open field test. Mifepristone decreased the total distance traveled and rearing, but it had no significant effect on horizontal beam breaks and stereotypies. The effects of mifepristone were not observed at the 24 h and 48 h time points, indicating that mifepristone did not have long-term effects on locomotor activity. Furthermore, these results indicate that mifepristone affects some behaviors in the open field test but not others. In a previous study, central (intracerebroventricular) administration of mifepristone decreased locomotor activity in rats [60]. Furthermore, both central and systemic mifepristone administration decreased locomotor activity induced by morphine, cocaine, and amphetamine in male Sprague Dawley, Fisher, and Wistar rats [60–62]. Central administration of mifepristone also decreases morphine-induced dopamine release in the nucleus accumbens (NAcc) and stress-induced dopamine release in the medial prefrontal cortex of rats [60, 63]. The reinforcing properties of nicotine are primarily mediated via the activation of dopamine neurons in the ventral tegmental area and the subsequent release of dopamine in the NAcc [64]. Noncontingent nicotine administration or nicotine self-administration enhances dopamine release in NAcc of rats [65, 66]. In addition, operant responding for food pellets induces dopamine release in the NAcc of rats [67]. Therefore, it is possible that the effects of mifepristone on nicotine and food intake, are at least in part, mediated via its effects on dopamine release in the NAcc. Considering the role of dopamine signaling in modulating reward function, the effect of mifepristone on dopamine release in the NAcc supports its potential use in treating nicotine addiction and obesity.

In order to gain insight into the effects of daily and intermittent nicotine intake on the cholinergic system, the effects of nicotine and mecamylamine on nicotine self-administration were investigated. Because the half-lifes (t1/2) of nicotine (t1/2, 1 h) and mecamylamine (t1/2, 1.2 h) in rats are relatively short, we also determined the time course effects of these drugs [68, 69]. Pre-treatment with nicotine led to a decrease in total nicotine intake in rats with daily and intermittent long access. The effect of nicotine was most pronounced during the first hour of access in both groups. The observation that pre-treatment with nicotine decreases nicotine intake is in line with the clinical studies that show that nicotine reduces the urge to smoke and nicotine intake in smokers [70–72]. In our study, pre-treatment with mecamylamine (2 mg/kg) did not affect total nicotine intake over the 6 h self-administration period in both daily long access (dependent) and intermittent long access (non-dependent) rats. Several previous studies have investigated the effects of mecamylamine on nicotine self-administration in rats. Patterson and Markou reported that mecamylamine (0.05 - 2 mg/kg) decreased nicotine intake in a combined group of nicotine-dependent daily short and long access rats (1 and 6 h, 7-days/week) and in non-dependent short access rats (1 h, 5 days/week)[50]. Other studies, including a study by Corrigall and Coen [41] with short (1 h) access rats and a study by DeNoble and Mele [73] with long (24 h) access rats, have also shown that mecamylamine (≤ 4 mg/kg) decreases nicotine intake. Of the aforementioned studies, only DeNoble and Mele specifically investigated the effects of mecamylamine in rats with long access to nicotine [73]. There are several differences between the present study, in which mecamyline did not decrease total nicotine intake, and the study by DeNoble and Mele, in which mecamylamine decreased total nicotine intake, that might explain the differential response to mecamylamine in these studies [73]. First, in our study, the rats had self-administered nicotine for about 7 weeks before the effects of mecamylamine on nicotine intake were investigated, while in the DeNobel and Mele study, the effects of mecamylamine were determined soon after nicotine intake stabilized [73]. Second, in the current study, the rats self-administered 0.06 mg/kg/inf of nicotine, while in the study by DeNoble and Mele, the rats self-administered 0.03 mg/kg/of nicotine [73]. Nicotine intake is higher in rats that self-administer 0.06 mg/kg/inf of nicotine than in rats that self-administer 0.03 mg/kg/inf of nicotine [40]. The prolonged self-administration period (7 weeks) and the high dose of nicotine (0.06 mg/kg/inf) might explain why mecamylaine did not reduce total nicotine intake over the 6 h period in this study.

We also investigated the time course effects of mecamylamine on nicotine intake over the 6 h nicotine self-administration period. Interestingly, although mecamylamine (2 mg/kg) did not affect the total 6 h nicotine intake in rats with daily access, it induced a large increase in nicotine intake during the first hour of access. This observation aligns with a previous study in which mecamylamine (0.75 and 1.5 mg/kg) increased nicotine intake during the first 3 h block of a 24 h self-administration session [73]. In the current study, we found that in the rats with intermittent long access, 2 mg/kg of mecamylamine had no effect on nicotine intake over the 6 h self-administration period and intake during the first hour of access. It is noteworthy that the response to mecamylamine of the rats with intermittent long access to nicotine falls between the responses of rats with daily short access and daily long access. Administration of 2 mg/kg of mecamylamine decreases first-hour nicotine intake in short-access (3 h) rats [74], had no effect on first-hour intake in the intermittent long-access (6 h) rats, and increased first-hour intake in the daily long-access (6 h) rats. These findings indicate that the effects of mecamylamine on nicotine intake are contingent upon the self-administration protocol. Mecamylamine decreases nicotine intake in non-dependent animals with short access to nicotine [41, 74]. In contrast, mecamylamine increased nicotine intake during the first hour of access in dependent rats with daily long access, while mecamylamine did not affect first-hour nicotine intake in non-dependent rats with intermittent long access. The effects of mecamylamine on nicotine intake in rats with daily long access are analogous to the effects of mecamylamine in regular smokers. Many studies have shown that mecamylamine increases smoking [75–78]. This phenomenon may be attributed to the fact that mecamylamine inhibits the psychoactive effects of nicotine, thereby leading to a compensatory increase in smoking [79, 80].

Mifepristone binds to both glucocorticoid receptors and progesterone receptors [19]. Although the effects of progesterone on drug intake have been thoroughly investigated, limited research has been conducted to investigate the effects of progesterone receptor blockade on drug intake [81]. A clinical study with male and female smokers demonstrated that progesterone administration reduces smoking urges [82]. Moreover, progesterone increased the aversive effects of intravenous nicotine and decreased the rewarding effects of intravenous nicotine [82]. However, sex differences were not evaluated in the aforementioned study. A study with male rats revealed that progesterone does not inhibit the reinstatement of nicotine-seeking [83]. At present, the effects of progesterone on the addictive properties of another stimulant, cocaine, are better understood than those of nicotine. Clinical studies suggest that progesterone mitigates the subjective effects of cocaine in females but not males [84]. Additionally, females with high progesterone levels have less stress and cue-induced cocaine craving than those with low levels [85]. Pre-clinical studies indicate that progesterone inhibits cocaine-induced conditioned place preference in female rats [86]. In the same study, castration did not affect cocaine-induced CPP, implying that gonadal hormones such as progesterone do not play a significant role in the rewarding effects of cocaine in male rats [86]. Furthermore, progesterone reduces cocaine-induced reinstatement of cocaine-seeking in female rats in estrus but not in other phases of the estrous cycle [87]. Additionally, it has been reported that estradiol administration enhances the acquisition of cocaine self-administration in ovariectomized female rats, an effect that is inhibited by progesterone [88]. These and other findings suggest that progesterone has a greater effect on the reinforcing efficacy of psychostimulants in females than males (for an extensive review on this topic see [89, 90]). Moreover, there is currently no evidence indicating that the blockade of progesterone receptors affects the reinforcing properties of nicotine. Therefore, it is unlikely that the effects of mifepristone on nicotine intake were mediated through progesterone receptor blockade.

In conclusion, this study investigated the effects of mifepristone on nicotine intake in rats with daily and intermittent long access to nicotine, as well as its effects on locomotor activity and operant responding for food pellets. Rats with daily access to nicotine developed nicotine dependence. Although the rats with intermittent access to nicotine had a higher level of nicotine intake per session, they did not become dependent. Mifepristone exhibited a greater reduction in nicotine intake in rats with intermittent access than in rats with daily access. Moreover, mifepristone decreased both the total distance traveled in the open field, and operant responding for food pellets. Additionally, mecamylamine increased nicotine intake during the first hour of access in rats with daily access, but not in rats with intermittent access. Pre-treatment with nicotine decreased nicotine intake in both rats with daily and intermittent access. These findings indicate that the history of nicotine intake affects the response to drugs that modulate nicotine intake. Moreover, glucocorticoid receptor blockade may offer a novel approach to reduce smoking in both regular and intermittent smokers.

## CRediT authorship contribution statement

**R. Chellian:** Conceptualization, Formal analysis, Investigation, Writing - Review & Editing, Visualization. **A. Behnood-Rod:** Investigation, Project administration. **A. Bruijnzeel:** Conceptualization, Formal analysis, Writing - Original Draft, Visualization, Supervision, Project administration, Funding acquisition.

