## Supplemental Materials for "Mifepristone decreases nicotine intake in dependent and non-dependent adult rats"

**Statistics**

**Baseline nicotine self-administration, 1 h sessions:** Baseline operant responding on the active and inactive levers and nicotine intake (3 sessions of 0.03 mg/kg/inf and 2 sessions of 0.06 mg/kg/inf) were analyzed with two-way ANOVAs, with access schedule as a between-subjects factor and time as a within-subjects factor (Fig. S1A-D). **Baseline nicotine self-administration, 6 h sessions:** Baseline operant responding on the active and inactive levers over the 29 daily self-administration sessions and 13 intermittent self-administration sessions were analyzed with two-way ANOVAs, with time as the within-subjects factor and lever as a between-subjects factor (Fig. S2A and S2B). Baseline nicotine intake over the 29 daily sessions and 13 intermittent sessions was analyzed with one-way ANOVAs, with time as the within-subjects factor (Fig. 2A and 2B). Operant responding on the active and inactive levers and nicotine intake in rats with daily and intermittent access over a period of 13 sessions was analyzed with two-way ANOVAs, with time as a within-subjects factor and access schedule as a between-subjects factor (Fig. S2C and S2D). Operant responding on the active and inactive levers and nicotine intake during the first and last session in rats with daily and intermittent access was analyzed with two-way ANOVAs, with session as a within-subjects factor and access schedule as a between-subjects factor (Fig. 2C, S2E, and S2F). The time course of nicotine intake in the first and the last session in rats with daily and intermittent access to nicotine was analyzed with three-way ANOVAs, with time point and session as within-subjects factors and access schedule as a between-subjects factor (Fig. 2D and 2E). **Mecamylamine-precipitated somatic withdrawal signs:** Somatic withdrawal scores were analyzed with a two-way ANOVA, with mecamylamine treatment as a within-subjects factor and access schedule as a between-subjects factor (Fig. 2F). **Mifepristone, mecamylamine, and nicotine treatments on nicotine self-administration in rats with daily and intermittent access:** The effects of mifepristone (90 min after), mecamylamine (10 min after), and nicotine (10 min after) treatments on operant responding on the active and inactive levers and nicotine intake were analyzed with two-way ANOVAs, with drug treatment as within-subjects factor and access schedule as between-subjects factor (Fig. 3A-C, 5A and 5C, S4A-D). The effects of mifepristone, mecamylamine, and nicotine treatments on the time course (1-6 h) of nicotine intake were analyzed with three-way ANOVAs, with time point and drug treatment as within-subjects factors and access schedule as a between-subjects factor (Fig. 4A-C, 5B and 5D). The effects of mifepristone, mecamylamine, and nicotine treatments (24 h and 48 h after) on operant responding on the active and inactive levers and nicotine intake in daily access rats were analyzed with one-way ANOVAs, with drug treatment as a within-subjects factor (Fig. S3A-C, S3E-G, S5A-L). The effects of mifepristone, mecamylamine, and nicotine treatments (24 h and 48 h after) on the time course (1-6 h) of nicotine intake in daily access rats were analyzed with two-way ANOVAs, with time point and drug treatment as a within-subjects factors (Fig. S3D and S3H, S6-D). **Mifepristone treatment on food responding and motor activity:** The effects of mifepristone on operant responding on the active and inactive levers, food pellets received, and open-field behavior were analyzed with one-way ANOVAs, with drug treatment as within-subjects factor (Fig. 6A-G, S7A-F, S8A-H). For all statistical analyses, significant effects in the ANOVA were followed by Bonferroni's posthoc tests to determine which groups differed from each other. P-values that were less or equal to 0.05 were considered significant. Significant main effects, interaction effects, and post hoc comparisons are reported in the Results section. Data were analyzed with SPSS Statistics version 29 and GraphPad Prism version 9.3.1.

**Table S1. Summary of ANOVA analyses**

| **Analysis** | **ANOVA Type** | **Within-Subjects Factors** | **Between-Subjects Factors** | **Figures** |
| --- | --- | --- | --- | --- |
| Baseline nicotine IVSA (1 h) | Two-way | Time | Access schedule | Fig. S1A-D |
| Baseline nicotine IVSA (6 h) | Two-way | Time | Lever | Fig. S2A, S2B |
| Baseline nicotine IVSA (daily/intermittent) | One-way | Time | - | Fig. 2A, 2B |
| Baseline nicotine IVSA (daily/intermittent access,13 sessions) | Two-way | Time | Access schedule | Fig. S2C, S2D |
| Baseline nicotine IVSA (first/last session) | Two-way | Session | Access schedule | Fig. 2C, S2E, S2F |
| Baseline nicotine IVSA (Time course, first/last session) | Three-way | Time point, Session | Access schedule | Fig. 2D |
| Mecamylamine-precipitated somatic withdrawal signs | Two-way | Mecamylamine treatment | Access schedule | Fig. 2F |
| Mifepristone, mecamylamine, nicotine treatments (nicotine IVSA, 10-90min) | Two-way | Drug treatment | Access schedule | Fig. 3A-C, 5A, 5C, S4A-D |
| Mifepristone, mecamylamine, nicotine treatments (nicotine IVSA, time course, 10-90min) | Three-way | Time point, Drug treatment | Access schedule | Fig. 4A-C, 5B, 5D |
| Mifepristone, mecamylamine, nicotine treatments (nicotine IVSA, 24h/48h) | One-way | Drug treatment | - | Fig. S3A-C, S3E-G, S5A-L |
| Mifepristone, mecamylamine, nicotine treatments (nicotine IVSA, time course, 24h/48h) | Two-way | Time point, Drug treatment | - | Fig. S3D, S3H, S6A-D |
| Mifepristone treatment on food responding & motor activity (nicotine self-admin, 90min, 24h/48h) | One-way | Drug treatment | - | Fig. 6A-G, S7A-F, S8A-H |

Abbreviations: IVSA, intravenous self-administration

**Results**

**Effects of mecamylamine on nicotine intake (24 and 48 h time-points)**

***24 and 48 h after mecamylamine treatment (daily group).*** Mecamylamine did not affect responding on the active and inactive lever and nicotine intake 24 h and 48 h later (24 h time point; Fig. S5A, Nicotine intake: Treatment F1,9 =4.31, NS; Fig. S5B, Active lever: Treatment F1,9 =3.746, NS; Fig. S5C, Inactive lever: Treatment F1,9 =1.159, NS; 48 h time point; Fig. S5D, Nicotine intake: Treatment F1,9 =0.469, NS; Fig. S5E, Active lever: Treatment F1,9 =0.1, NS; Fig. S5F, Inactive lever: Treatment F1,9 =0.091, NS). ***Time course analysis, 24 and 48 h after mecamylamine treatment (daily group).*** Mecamylamine did not affect the time course of nicotine intake 24 h (Fig. S6A, Nicotine intake: Treatment F1,9 =4.31, NS; Time point F5,45 =39.172, P < 0.001; Treatment x Time point F5,45=1.312, NS) or 48 h (Fig. S6B, Nicotine intake: Treatment F1,9 =0.469, NS; Time point F5,45 =20.373, P < 0.001; Treatment x Time point F5,45=1.305, NS) later.

**Effects of nicotine pretreatment on nicotine intake (24 and 48 h time-points)**

***24 and 48 h after nicotine treatment (daily group).*** Nicotine treatment did not affect nicotine intake and responding on the active lever and inactive lever 24 h and 48 h later in the rats with daily access to nicotine (24 h time point; Fig. S5G, Nicotine intake: Treatment F1,9 =2.983, NS; Fig. S5H, Active lever: Treatment F1,9 =2.903, NS; Fig. S5I, Inactive lever: Treatment F1,9 =2.78, NS; 48 h time point; Fig. S5J, Nicotine intake: Treatment F1,9 =0.078, NS; Fig. S5K, Active lever: Treatment F1,9 =0.108, NS; Fig. S5L, Inactive lever: Treatment F1,9 =1.256, NS). ***Time course analysis, 24 and 48 h after nicotine treatment.*** Nicotine treatment did not affect the time course of nicotine intake 24 h (Fig. S6C, Nicotine intake: Treatment F1,9 =2.983, NS; Time point F5,45 =44.125, P < 0.001; Treatment x Time point F5,45=0.566, NS) or 48 h later (Fig. S6D, Nicotine intake: Treatment F1,9 =0.078, NS; Time point F5,45 =24.775, P < 0.001; Treatment x Time point F5,45=0.303, NS).

**Effect of mifepristone on operant responding for food (24 and 48 h time-points)**

***24 h after mifepristone treatment*.** Treatment with mifepristone did not affect food intake, responding on the active lever and inactive lever 24 h after treatment (Fig. S7A, Food intake: Treatment F3,27 =0.766, NS; Fig. S7B, Active lever: Treatment F3,27 =0.979, NS; Fig. S7C, Inactive lever: Treatment F3,27 =0.802, NS). ***48 h after mifepristone treatment*.** Furthermore, mifepristone did not affect food intake, responding on the active lever and inactive lever 48 h after treatment (Fig. S7D; Food intake: Treatment F3,27 =0.709, NS; Fig. S7E, Active lever: Treatment F3,27 =0.873, NS; Fig. S7F, Inactive lever: Treatment F3,27 =1.231, NS).

**Effect of mifepristone on locomotor activity (24 and 48 h time-points)**

***24 h after mifepristone treatment*.** Treatment with mifepristone did not affect the total distance traveled, horizontal beam breaks, vertical beam breaks, and stereotypies (Fig. S8A, Total distance: Treatment F3,27 =1.213, NS; Fig. S8B, Horizontal beam breaks: Treatment F3,27 =0.646, NS; Fig. S8C, Vertical beam breaks: Treatment F3,27 =0.264, NS; Fig. S8D, Stereotypies: Treatment F3,27 =0.583, NS*).* ***48 h after mifepristone treatment*.** Treatment with mifepristone did not affect the total distance traveled, horizontal beam breaks, vertical beam breaks, and stereotypies (Fig. S8E, Total distance: Treatment F3,27 =0.208, NS; Fig. S8F, Horizontal beam breaks: Treatment F3,27 =0.037, NS; Fig. S8G, Vertical beam breaks: Treatment F3,27 =0.776, NS; Fig. S8H, Stereotypies: Treatment F3,27 =0.224, NS).

**Tables**

**Table S2. Effects of mifepristone, mecamylamine, and nicotine treatment on nicotine Intake and time course effects**

| **Drug** | **Effect on Total Nicotine Intake** | **Time Course Effect (Daily Access)** | **Time Course Effect (Intermittent Access)** | **Figures** |
| --- | --- | --- | --- | --- |
| Mifepristone | Decrease in daily and intermittent access rats | Decrease for 1 h | Decrease for 2 h | Fig. 3A, 4A, 4B |
| Mecamylamine | No effect in daily and intermittent access rats | Increase for 1 h | No effect | Fig. 5A, 5B |
| Nicotine | Decrease in daily and intermittent access rats | Decrease for 1 h | Decrease for 1 h | Fig. 5C, 5D |

**Figures**

**Figure S1**

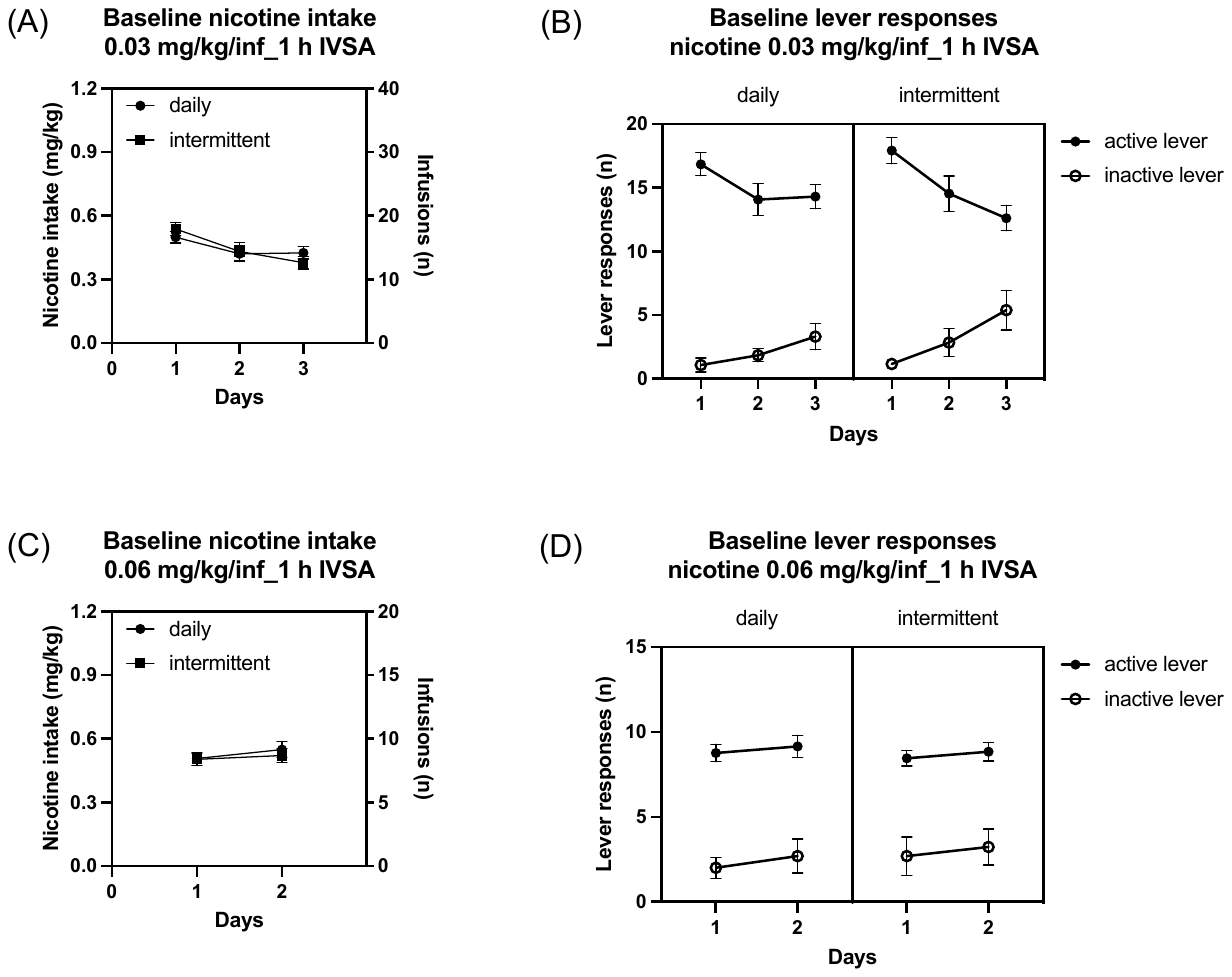

**Figure S1. Baseline nicotine intake in rats.** At the onset of the self-administration study, the rats self-administered 0.03 mg/kg/inf of nicotine for three days (A, nicotine intake; B, lever responses) and then 0.06 mg/kg/inf of nicotine for two days in 1 h sessions (C, nicotine intake; D, lever responses). During the first three sessions, nicotine intake and responding on the active lever decreased, and responding on the inactive lever increased (A, B). During the following two sessions, there was no change in nicotine intake and responding on the active and inactive lever (C, D). Daily n=13, Intermittent, n=13. Data are expressed as means ± SEM.

**Figure S2**

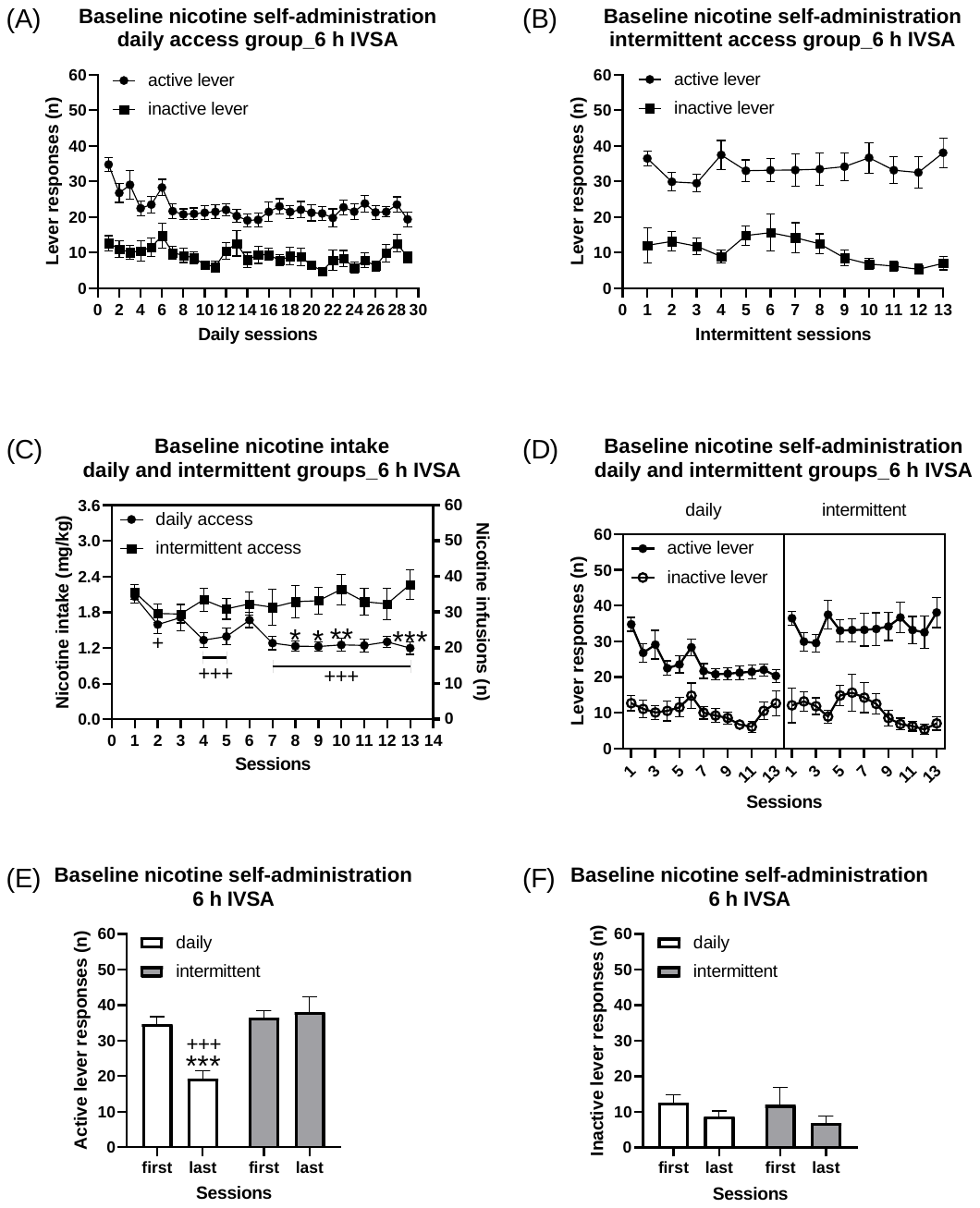

**Figure S2. Baseline nicotine intake in rats with daily and intermittent access.** The rats with daily access self-administered nicotine for 29 sessions and the rats with intermittent access self-administered nicotine for 13 sessions. All the sessions lasted 6 h and the rats self-administered 0.06 mg/kg/inf of nicotine. The figures depict responding on the active and inactive lever in rats with daily (A) and intermittent access to nicotine (B). The figures show nicotine intake (C) and lever pressing (D) in rats with daily and intermittent access during the first 13 sessions. The rats with intermittent access had a higher level of nicotine intake and more active lever presses than rats with daily access (C, D). The access schedule did not affect responding on the inactive lever (C). The figures also depict responding on the active (E) and inactive lever (F) during the first and the last nicotine self-administration session in rats with daily and intermittent access. Plus signs indicate lower nicotine intake compared to nicotine intake of the same group on Day 1 (C) and fewer active lever responses compared to rats with intermittent access during the last session (E). Asterisks indicate lower nicotine intake in rats with daily access than in rats with intermittent access (C) and lower active lever responses compared to the first session of rats in the same group (E). *, + P<0.05; ** P<0.01; +++, *** P<0.001. Daily n=13, Intermittent, n=13. Data are expressed as means ± SEM.

**Figure S3**

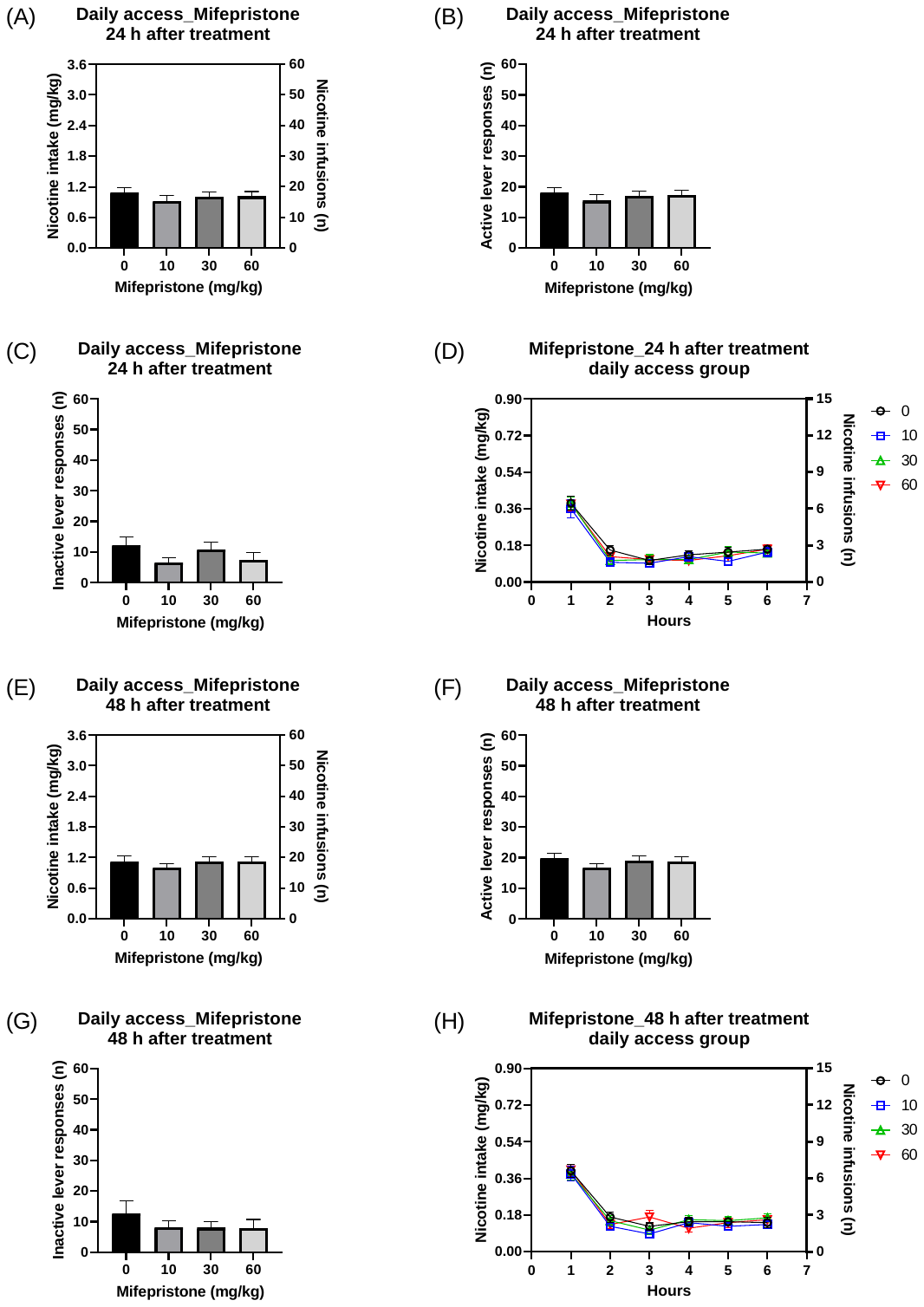

**Figure S3. Mifepristone does not have a lasting effect on nicotine intake.** The effects of mifepristone on nicotine self-administration and the time course of nicotine intake was investigated 24 h and 48 h after treatment. Mifepristone did not affect nicotine intake (A, 24 h; E, 48 h), active lever responses (B, 24 h; F, 48 h), and inactive lever responses (C, 24 h; G, 48 h) 24 h and 48 h after treatment. Mifepristone also did not affect the time course of nicotine intake 24 h and 48 h after treatment (D, H). Daily n=13, Intermittent, n=13. Data are expressed as means ± SEM.

**Figure S4**

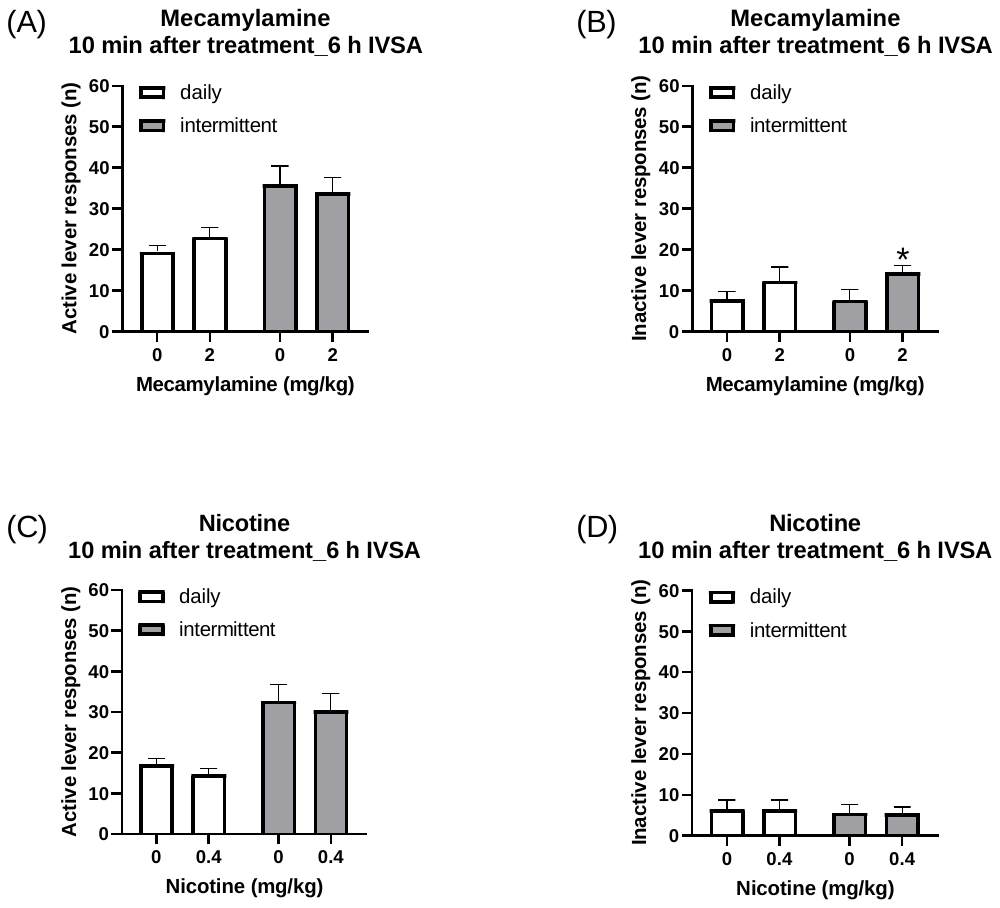

**Figure S4. Pre-treatment with mecamylamine and nicotine on lever responses.** Mecamylamine did not affect responding on the active lever (A) but increased responding on the inactive lever (B) in rats with daily and intermittent access to nicotine. Pre-treatment with nicotine decreased responding on the active lever but did not affect responding on the inactive lever in rats with daily and intermittent access to nicotine (C, D). Asterisks indicate more inactive lever responses in rats treated with 2 mg/kg of mecamylamine than in vehicle-treated rats with intermittent access. * P<0.05. Daily n=10, Intermittent, n=13. Data are expressed as means ± SEM.

**Figure S5**

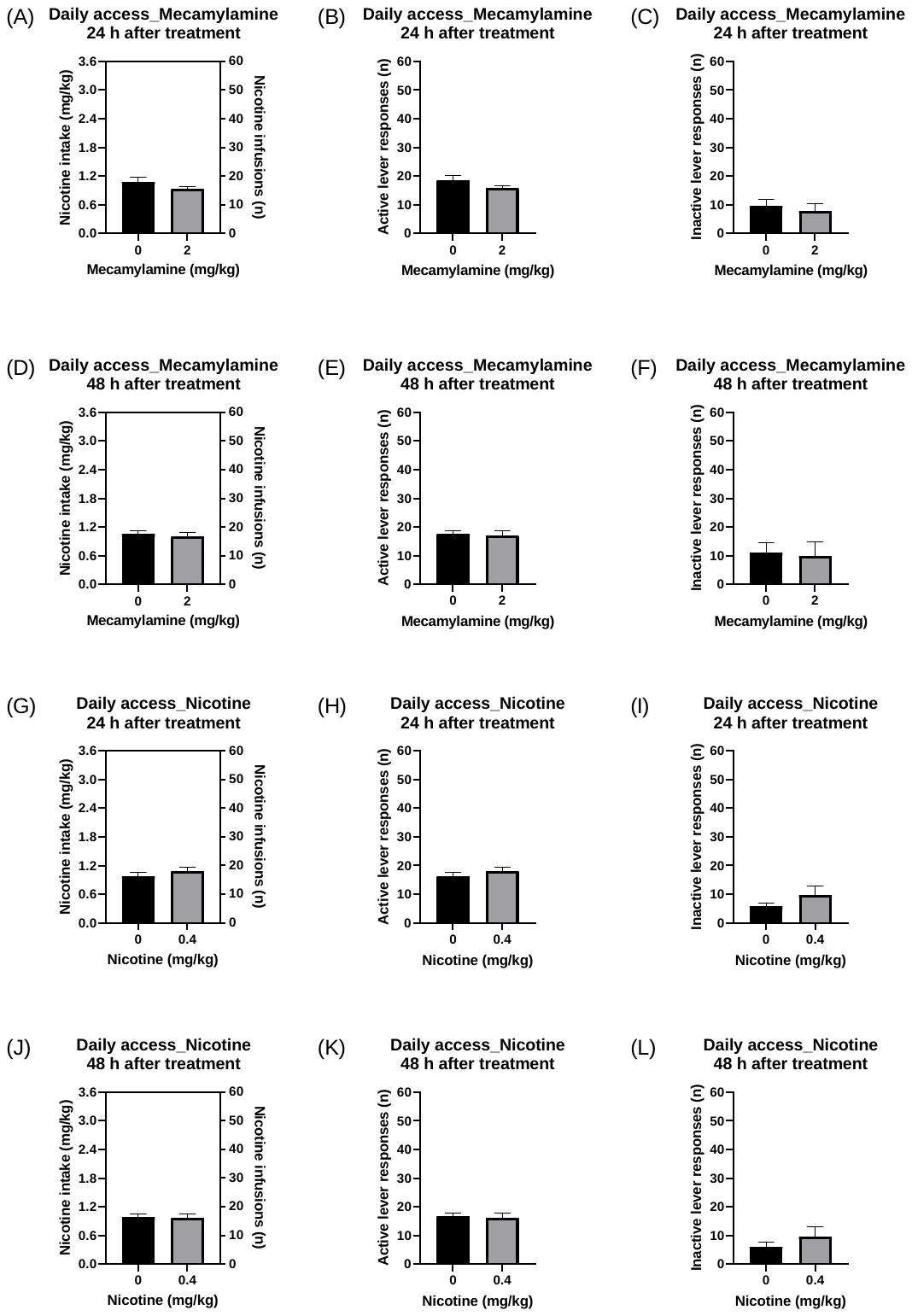

**Figure S5. Mecamylamine and nicotine treatment does not have a lasting effect on nicotine intake.** The effects of mecamylamine and nicotine treatment on nicotine self-administration was investigated 24 h and 48 h after treatment. Mecamylamine treatment did not affect nicotine intake (A, 24 h; D, 48 h), active lever responses (B, 24 h; E, 48 h), and inactive lever responses (C, 24; F, 48 h) 24 h and 48 h after treatment. Nicotine treatment did not affect nicotine intake (G, 24 h; J, 48 h), active lever responses (H, 24 h; K, 48 h), and inactive lever responses (I, 24 h; L, 48 h) 24 h and 48 h after treatment. Daily n=10, Intermittent, n=13. Data are expressed as means ± SEM.

**Figure S6**

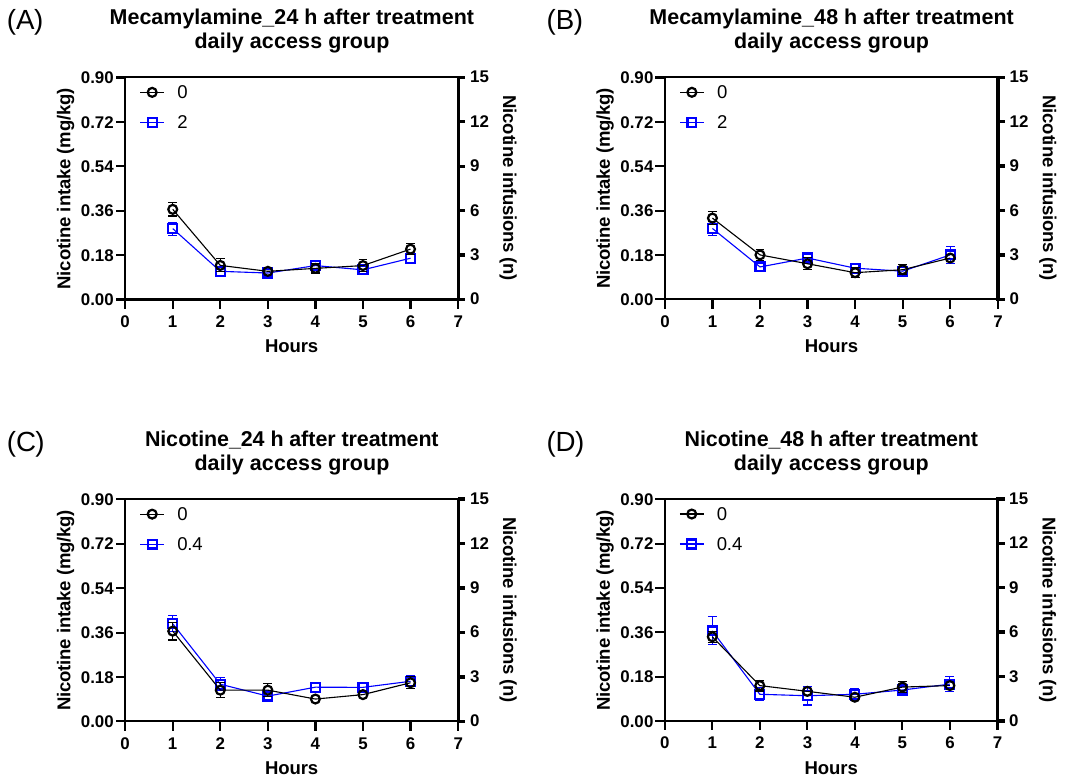

**Figure S6. Mecamylamine and nicotine treatment does not have a lasting effect on the time course of nicotine intake.** The effects of mecamylamine and nicotine treatment on the time course of nicotine intake were investigated 24 h and 48 h after treatment. Mecamylamine (A, 24 h; B, 48 h) and nicotine (C, 24 h; D, 48 h) treatment did not affect the time course of nicotine intake 24 h and 48 h after treatment. Daily n= 10, Intermittent, n=13. Data are expressed as means ± SEM.

**Figure S7**

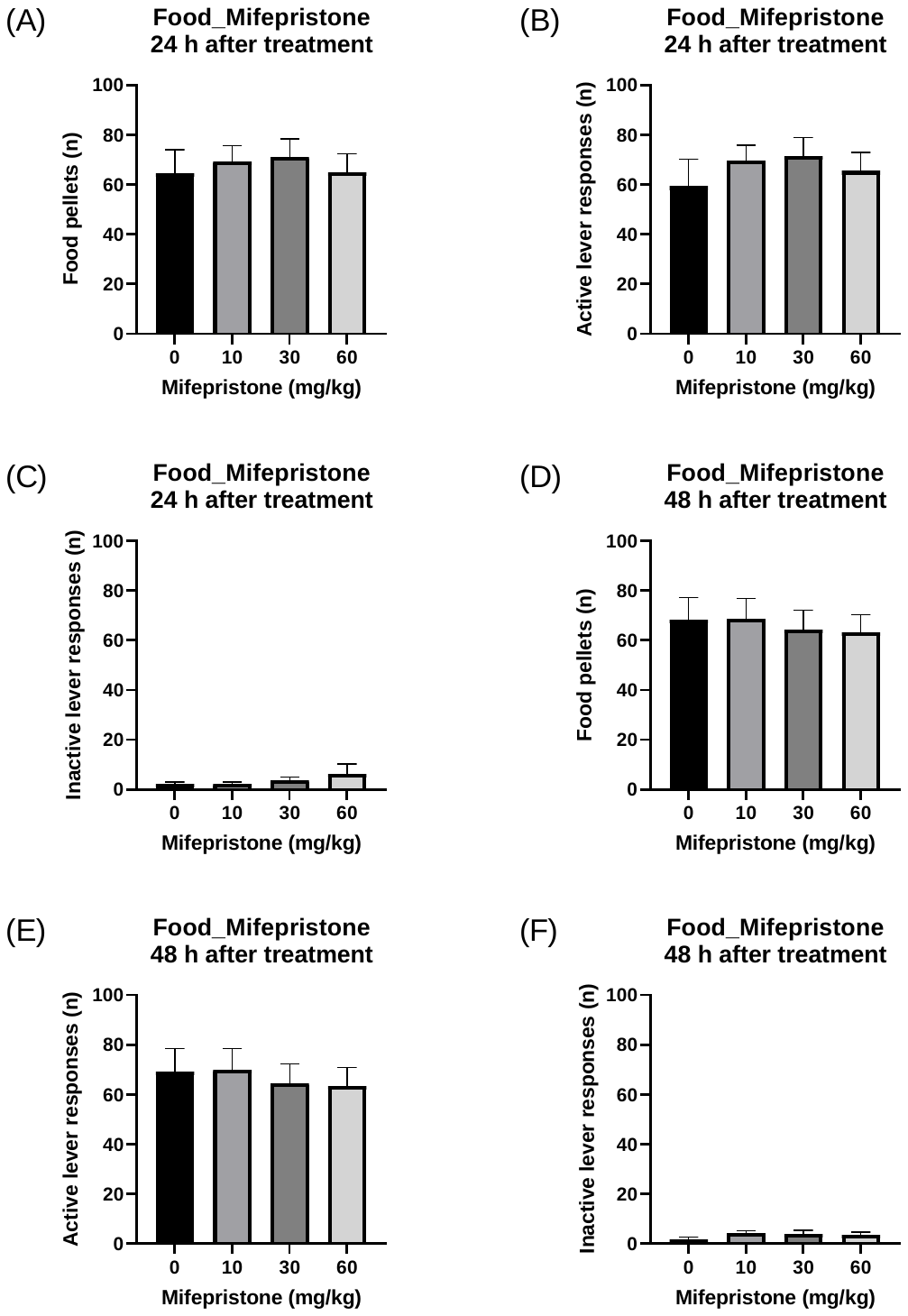

**Figure S7. Mifepristone treatment does not have a lasting effect on operant responding for food.** The effects of mifepristone on the number of food pellets received (A, 24 h; D, 48 h), active lever responses (B, 24 h; E, 48 h), and inactive lever responses (C, 24 h; F, 48 h) were evaluated. Mifepristone did not affect the number of food pellets, active lever responses, and inactive lever responses 24 h and 48 h after treatment. N=10. Data are expressed as means ± SEM.

**Figure S8**

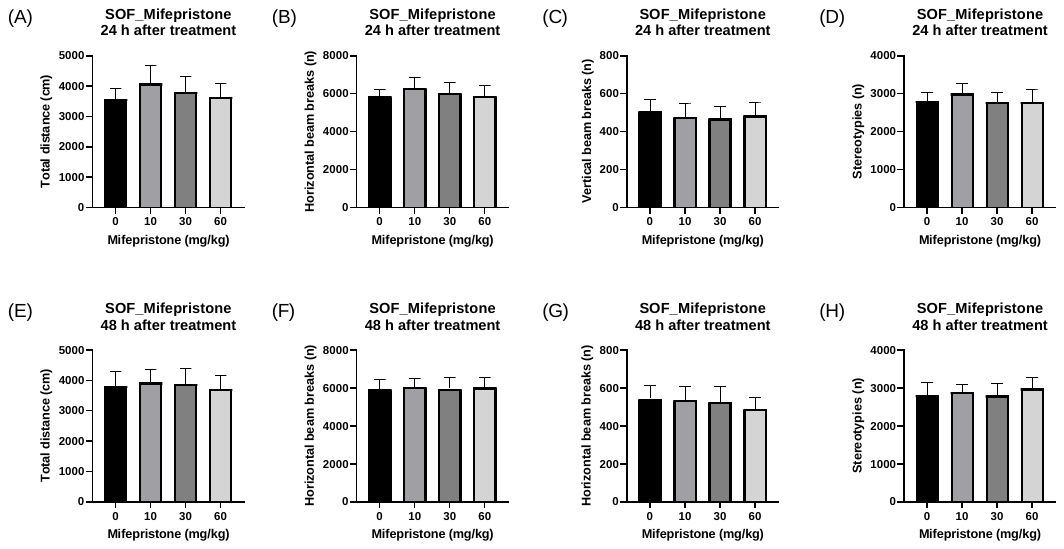

**Figure S8. Mifepristone treatment does not have a lasting effect on locomotor activity.** The effects of mifepristone on the total distance traveled (A, 24 h; E, 48 h), horizontal beam breaks (B, 24 h; F, 48 h), vertical beam breaks (C, 24 h; G, 48 h), and stereotypies (D, 24 h; H, 48 h) were evaluated. Mifepristone did not affect the total distance traveled, horizontal beam breaks, vertical beam breaks, and stereotypies 24 h and 48 h after treatment. N=10. Data are expressed as means ± SEM.
